# Presence of the Hmq system and production of 4-hydroxy-3-methyl-2-alkylquinolines is heterogeneously distributed between *Burkholderia cepacia* complex species and more prevalent among environmental than clinical isolates

**DOI:** 10.1101/2020.08.27.269993

**Authors:** Pauline M.L. Coulon, James E.A. Zlosnik, Eric Déziel

**Affiliations:** Centre Armand-Frappier Santé Biotechnologie, Institut National de la Recherche Scientifique (INRS), Laval, QC, Canada; Canadian Burkholderia cepacia Complex Research and Referral Repository, University of British Columbia, Vancouver, BC, Canada

**Keywords:** *hmqABCDEFG* operon, 4-hydroxy-2-alkylquinolines (HAQs), *pqsABCDE*, *Pseudomonas* Quinolone Signal (PQS), quorum sensing (QS)

## Abstract

Some *Burkholderia cepacia* complex (Bcc) strains have been reported to produce 4-hydroxy-3-methyl-2-alkylquinolines (HMAQs), analogous to the 4-hydroxy-2-alkylquinolines of *Pseudomonas aeruginosa*. Using *in silico* analyses, we previously showed that the *hmqABCDEFG* operon, which encodes enzymes involved in the biosynthesis of HMAQs, is carried by about one-third of Bcc strains, with considerable inter- and intra-species variability. In the present study, we investigated by PCR, using consensus primers, the distribution of *hmqABCDEFG* in a collection of 313 Bcc strains (222 of clinical and 91 of environmental origins) - belonging to 18 Bcc species. We confirmed that the distribution is species-specific, although not all strains within a species carry the *hmqABCDEFG* operon. Among the 30% of strains bearing the *hmqABCDEFG* operon, we measured the total HMAQs production and showed that 90% of environmental isolates and 68% of clinically isolated Bcc produce detectable levels of HMAQs when cultured in TSB medium. For the strains having the *hmqABCDEFG* operon but not producing HMAQs, we studied the transcription and showed that none expressed the *hmqA* gene under the specified culture conditions. Interestingly, the *hmqABCDEFG* operon is more prevalent among plant root environment species (e.g. *B. ambifaria, B. cepacia*) and absent in species commonly found in chronically colonized individuals with cystic fibrosis (e.g. *B. cenocepacia, B. multivorans*), suggesting that the Hmq system could play a role in niche adaptation by influencing rhizosphere microbial community and could have been lost through evolution. Understanding the Hmq system and its regulation will provide clues concerning the production of HMAQs and their functions in Bcc.

## Introduction

The environmental species of *Burkholderia* can be divided into two phylogenic groups: (1) pathogenic species and (2) plant-beneficial species (Eberl and Vandamme, 2016). Based on this still controversial separation (Vandamme *et al*., 2017), the latter group was reclassified as *Paraburkholderia*, based on lower %GC and lack of virulence in *Caenorhabditis elegans* (Angus *et al*., 2014; Sawana *et al*., 2014). The pathogenic *Burkholderia* group comprises (1) plant pathogens (e.g.: *Burkholderia andropogonis* causing leaf streak on Sorghum (Ramundo and Claflin, 2005) and *B. glumae* causing bacterial panicle blight on rice (Azegami *et al*., 1985; Nandakumar *et al*., 2009; Ham *et al*., 2010)); (2) the “*pseudomallei*” group, comprised on *B. pseudomallei* (the causative agent of melioidosis), *B. thailandensis* (avirulent model) and *B. mallei* (causing glanders in equids) species (Howe, 1950; Smith *et al*., 1987; Chaowagul *et al*., 1989; Sandford, 1990) - and finally (3) the *Burkholderia cepacia* complex (Bcc), comprising at least 26 different species [(Bach *et al*., 2017; Martina *et al*., 2017; Weber and King, 2017); reviewed by (Eberl and Vandamme, 2016)], most considered opportunistic pathogens.

Bcc bacteria have been used in (1) agriculture for biocontrol of phytopathogens and plant growth-promoting properties [e.g.: pea protection by *B. ambifaria* against *Pythium* and *Aphanomyces* (Parke, 1991; Mullins *et al*., 2019)] and (2) bioremediation [e.g.: *B. vietnamiensis* with its trichloroethylene degradation abilities (Gillis *et al*., 1995; O’Sullivan and Mahenthiralingam, 2005)]; reviewed in Vial *et al*. (2011). Bcc bacteria are also well known for secondary metabolite production, including antibiotics [reviewed by Depoorter *et al*., (2016)].

However, in the 1980s, Bcc opportunistic pathogens have emerged as a serious issue among certain immunocompromised individuals (e.g. with the chronic granulomatous disease) and people with cystic fibrosis (CF) causing the ‘cepacia syndrome’, pushing authorities to prohibit their use in biotechnological applications. It is now generally accepted that their high transmissibility and intrinsic resistance to clinically relevant antibiotics makes them particularly problematic (Gilligan, 1991; Speert *et al*., 1994; Govan and Deretic, 1996).

Cell-to-cell communication mechanisms, e.g. quorum sensing (QS), act by (1) controlling gene transcription at the population level (Fuqua and Greenberg, 1975), (2) promoting colonization, and (3) optimizing interaction with hosts and increasing resistance to stress (Stewart and Costerton, 2001; Juhas *et al*., 2005). CepR/CepI is the primary QS system in Bcc species. The CepI synthase produces the autoinducer ligand N-octanoyl-homoserine lactone (C_8_-HSL) which interacts with the transcriptional regulator CepR to activate the transcription of several target genes, such as genes involved in the production of pyrrolnitrin (tryptophan halogenase (*prnA*); BAMB_RS23660), enacyloxins (LuxR family transcriptional regulator; BAMB_RS29445), and occidiofungins (amino acid adenylation domain-containing protein; BAMB_RS32210) (McKenney *et al*., 1995; Lewenza *et al*., 1999; Lewenza and Sokol, 2001; Chapalain *et al*., 2013). Depending on the Bcc species, at least two other Cep-like systems may be present, synthesizing different acyl-homoserine lactones (AHLs) as ligands and regulating each other (Choudhary *et al*., 2013).

The bacterium *Pseudomonas aeruginosa* carries a distinct QS system whose ligands are not AHLs but instead 4-hydroxy-2-alkylquinolines (HAQs), such as the *Pseudomonas* Quinolone Signal (PQS) and 4-hydroxy-2-heptylquinoline (HHQ) (Pesci *et al*., 1999; Déziel *et al*., 2004; Heeb *et al*., 2011). Interestingly, some strains of Bcc (*B. ambifaria* and *B. cepacia*), as well as *B. pseudomallei* and *B. thailandensis*, produce homologous molecules, that we call 4-hydroxy-3-methyl-2-alkylquinolines (Diggle *et al*., 2006; Vial *et al*., 2008; Ritzmann *et al*., 2019), that are synthesized by enzymes encoded by the *hmqABCDEFG* operon (Vial *et al*., 2008). In contrast with *P. aeruginosa* HAQs, the main HMAQs produced by *Burkholderia* bear a methyl group at the 3’ position and an unsaturation of the alkyl side chain. An increasing number of Bcc strains are being reported to produce some HMAQs congeners (Mori *et al*., 2007; Vial *et al*., 2008; Kilani-Feki *et al*., 2011, 2012; Mahenthiralingam *et al*., 2011; Li *et al*., 2018), but this remains mostly anecdotal. In contrast with the HAQ/PQS system of *P. aeruginosa*, the *Burkholderia* Hmq system does not appear to be a QS system *per se*, although we have shown that it is closely interrelated with the Cep system in *B. ambifaria* HSJ1 and the three Bta QS systems in *B. thailandensis* E264 (Vial *et al*., 2008; Chapalain *et al*., 2017; Le Guillouzer, 2018).

Functions of HHQ and PQS in *P. aeruginosa* as QS inducers, immunomodulators, antimicrobials have been described (Lin *et al*., 2018). Several studies report novel molecules belonging to the HAQ family and various bacterial species having antimicrobial activities (Wratten *et al*., 1977; Hamasaki *et al*., 2000; Whalen *et al*., 2014; Meyer *et al*., 2017; Dow *et al*., 2019). Only a few functions of HMAQs are known, apart from intra and interspecies QS signal (Vial *et al*., 2008; Chapalain *et al*., 2017; Le Guillouzer, 2018), especially as antimicrobials -having a lower activity than antibiotics -however, their main function remains enigmatic (Mori *et al*., 2007; Kilani-Feki *et al*., 2011, 2012; Mahenthiralingam *et al*., 2011; Li *et al*., 2018; Piochon *et al*., 2020). Nonetheless, given the demonstrated role of HAQs and PQS in *P. aeruginosa*, HMAQs may also play a role in the virulence and pathogenicity of opportunistic *Burkholderia* pathogens (Vial *et al*., 2008, 2009; Chapalain *et al*., 2017). We have previously characterized (Vial *et al*. (2008, 2009)) a series of clinical *B. ambifaria* strains able to produce HMAQs and that phenotypic variant of these *B. ambifaria* strains had lost their ability to produce several secondary metabolites, including HMAQs, similar a set of environmental isolates. Therefore, to better understand these metabolites, we posited the hypotheses that (1) the *hmqABCDEFG* operon is more frequently found among clinical Bcc strains and that (2) the clinical Bcc strains produce higher concentrations of HMAQs than environmental ones. Our previous bioinformatic study of the distribution of the *hmqABCDEFG* operon in the Bcc, based on available 1,257 whole-genome sequences belonging to 21 Bcc species, showed that strains belonging to the *B. ambifaria, B. cepacia, B. contaminans B. pyrrocinia, B. stagnalis. B. territorii*, and *B. ubonensis* species carry the *hmqABCDEFG* operon, but not all strains within a species, while *B. anthina, B. arboris B. cenocepacia, B. diffusa, B. latens, B. metallica, B. multivorans, B. pseudomultivorans, B. seminalis, B*.*stabilis and B. vietnamiensis* - species mainly found in clinical cases - are lacking *hmqABCDEFG* operon (Coulon *et al*., 2019). To experimentally validate our *in silico* study and verify the ability of Bcc isolates carrying the *hmqABCDEFG* operon to actually produce HMAQs, a collection of 313 Bcc strains, comprising 222 clinical and 91 environmental isolates belonging to 18 different Bcc species, was analyzed to first determine the presence of the *hmqABCDEFG* operon in their genome. We then directly determined, by liquid chromatography coupled to mass spectrometry (LC/MS) analyses, the ability of all the strains bearing the *hmqABCDEFG* operon to produce HMAQs. Finally, we verified the expression of *hmqA* in Bcc strains having the *hmqABCDEFG* and not producing HMAQs to investigate this lack of HMAQ production. Our data confirm that the Hmq system is heterogeneously distributed between Bcc species, with high prevalence in some species (e.g. *B. cepacia*) and near absence in other (e.g. *B. cenocepacia* and *B. multivorans*). Globally, higher frequency among strains of environmental origins vs clinical isolates suggests that the *hmqABCDEFG* operon and HMAQ production could play a role in Bcc niche adaptation.

## Results

### The *hmqABCDEFG* operon is heterogeneously distributed across and within Bcc species

We previously examined 1,257 whole-genome sequences belonging to 21 different Bcc species to assess the distribution of the *hmqABCDEFG* operon (Coulon *et al*., 2019). We found that at least one sequenced strain belonging to 7 out of 21 species carries the *hmqABCDEFG* operon (*B. ambifaria, B. cepacia, B. contaminans B. pyrrocinia, B. stagnalis. B. territorii*, and *B. ubonensis*); one striking initial finding was that prevalence of the *hmqABCDEFG* operon within a species appeared highly variable (Coulon *et al*., 2019). Here, to validate our *in silico* analyses of the distribution of the *hmqABCDEFG* operon based on homology and orthology (Coulon *et al*., 2019) and to globally determine the ability of Bcc to produce HMAQs, we screened a collection of 313 Bcc strains (222 of clinical and 91 of environmental origins; listed in **Table S1**), belonging to 18 Bcc species: *B. ambifaria, B. anthina, B. arboris, B. cenocepacia, B. cepacia, B. contaminans, B. diffusa, B. dolosa, B. lata, B. metallica, B. multivorans, B. pyrrocinia, B. seminalis, B. stabilis, B. stagnalis. B. territorii, B. ubonensis, B. vietnamiensis*, plus a few more classified in the ‘other Bcc’ group (PubMLST database; https://pubmlst.org/bcc/info/protocol.shtml) - for the presence of *hmqABCDEFG* by PCR using consensus primers targeting *hmqA* and *hmqG*. We had previously determined that the presence of a *hmqG* orthologue correlates with the presence of a complete *hmqABCDEFG* operon (Coulon *et al*., 2019). Here, we found that 30% of the tested strains possess an *hmqABCDEFG* operon, including 53% of environmental but only 21% of clinical strains (**Figure 1A**). Among the 18 different species investigated, 14 comprise at least one strain carrying the operon (**Figure 1B**).

**Figure 1.**
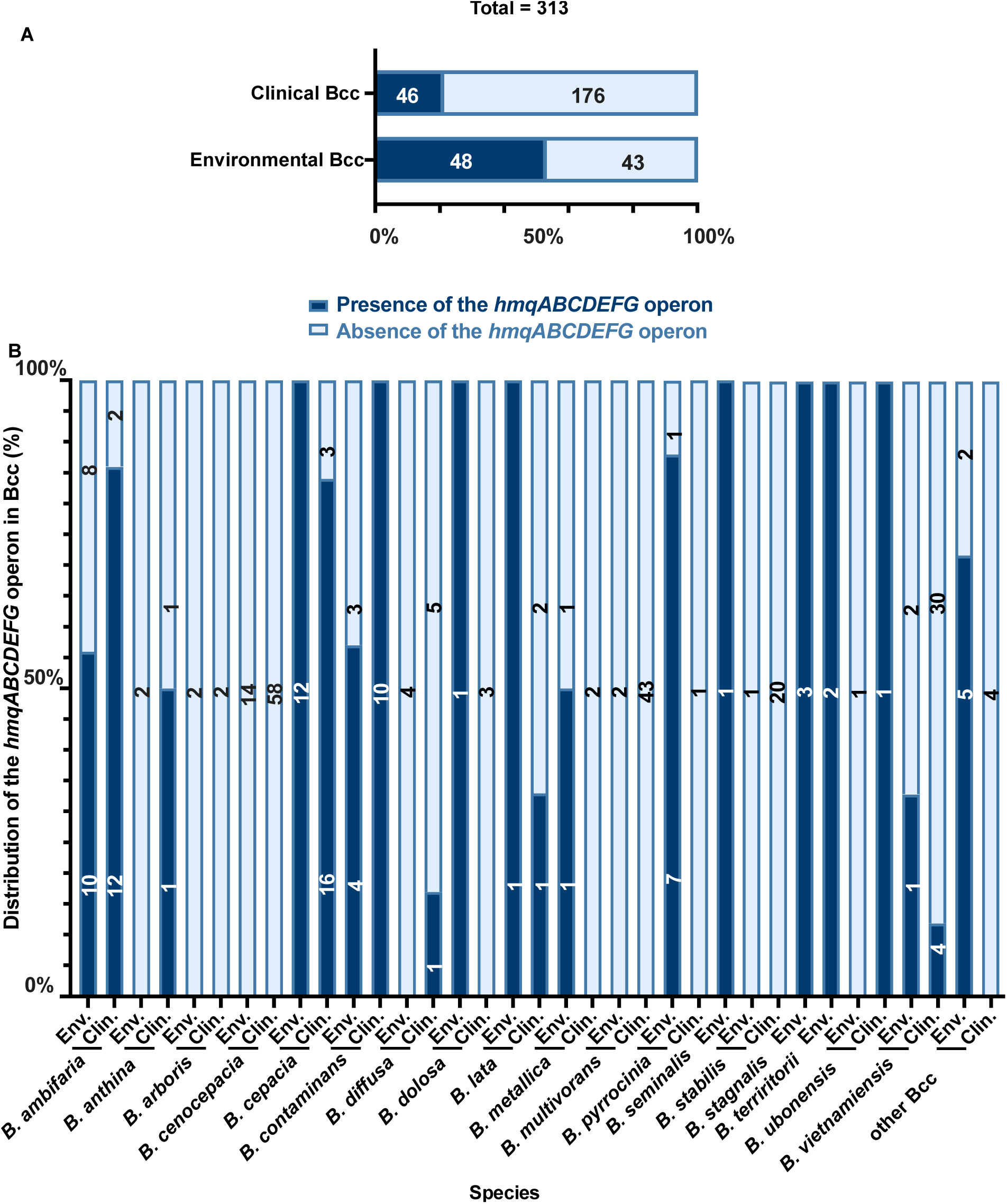
Bcc strains screening for the *hmqABCDEFG* operon presence in their genome. (A) The distribution of environmental and clinical Bcc strains investigated in this study (B) Distribution of the *hmqABCDEFG* operon within tested Bcc species.

The *hmqABCDEFG* operon is more prevalent among clinical strains for species *B. ambifaria, B. anthina, B. contaminans, B. diffusa, B. ubonensis* and *B. vietnamiensis*. However, it is more prevalent among environmental strains for *B. contaminans, B. cepacia, B. dolosa, B. lata, B. metallica, B. pyrrocinia*, and the ‘other Bcc’ group. Clinical *B. seminalis* and environmental *B. stagnalis* and *B. territorii* species carry *hmqABCDEFG*.

We found that isolates of *B. dolosa, B. anthina*, and *B. vietnamiensis* carry the *hmqABCDEFG* operon, which was not predicted in our previous analysis of available genomic data (Coulon *et al*., 2019), we confirmed here our PCR results by sequencing of the amplicons using primers listed in **Table S2**.

### Neither the phylogeny of Bcc species nor the loss of the third chromosome explain the distribution of the *hmqABCDEFG* operon

Since not all Bcc species carry the *hmqABCDEFG* operon, we asked whether the distribution of the operon could be related to the phylogenic distribution of Bcc species. Based on MultiLocus Sequence Typing (MLST) sequences, we found that *B. ambifaria, B. cepacia, B. contaminans, B. pyrrocinia* and *B. stagnalis* species *in* which *hmqABCDEFG* is the most prevalent, are not clustered (**Figure 2**). The same was observed for *B. cenocepacia* and *B. multivorans* which both do not possess the *hmqABCDEFG* operon (**Figure 2**).

**Figure 2.**
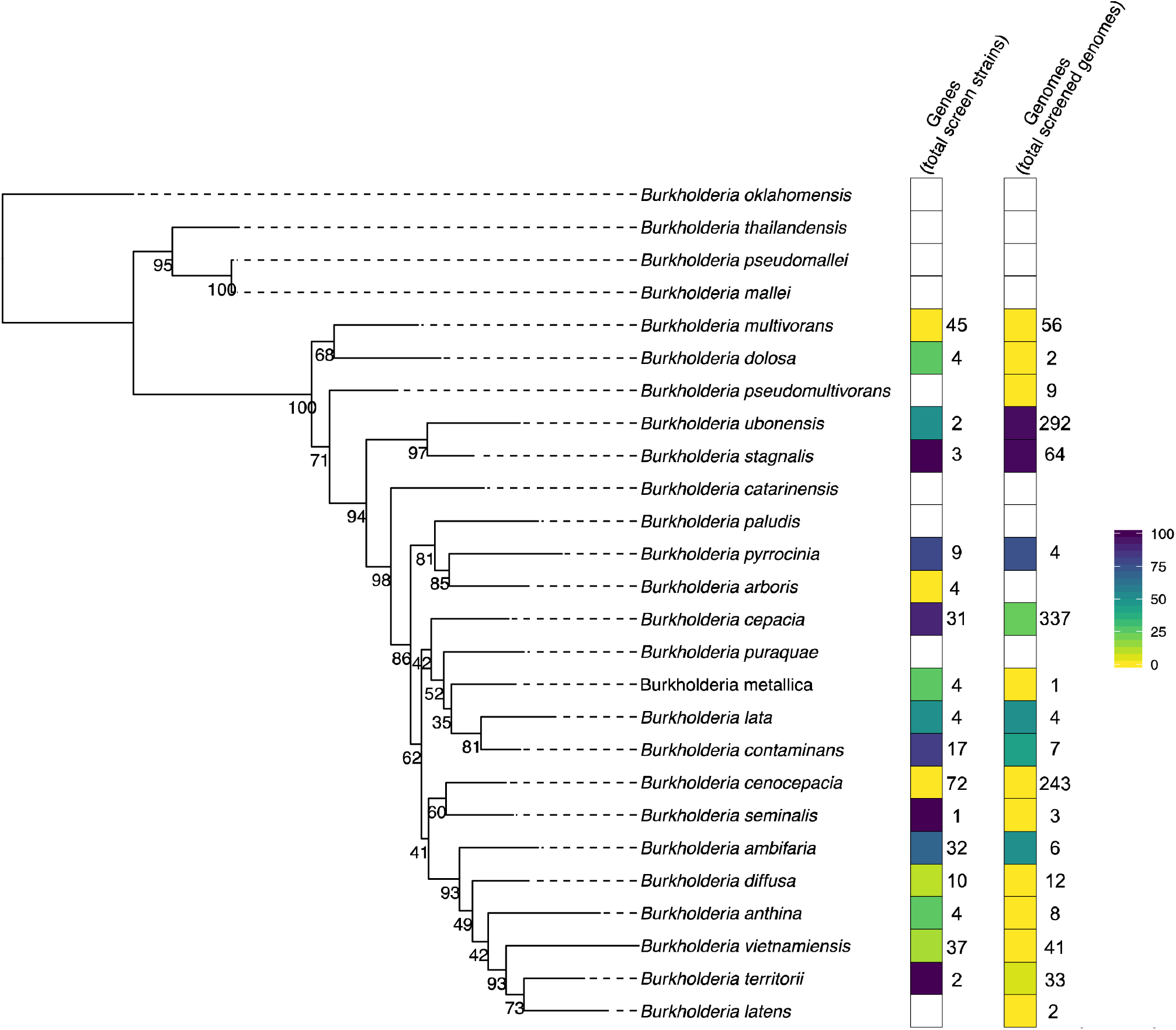
Phylogeny of the Bcc species based on MLST genes (*atpD, gltB, gyrB, recA, lepA, phaC* and *trpB*). The tree was generated by RAxML and is the concatenated MLST sequences (typically of the type strain for a given species) using GTRGAMMA model and 1000 bootstraps. The branches are labelled where bootstrap values are >50%. The presence of the *hmqABCDEFG* operon is correlated between the bioinformatic (Coulon *et al*., 2019) and PCR methods (Kendall rank test with a p-value of 0.01958, inferior to 5%).

Bcc bacteria are known to lose their third chromosome (c3) which is a virulence mega-plasmid containing a few core genes (Agnoli *et al*., 2011; diCenzo *et al*., 2019). The *hmqABCDEFG* operon being generally located on the c3 replicon, we investigated whether the absence of the *hmqABCDEFG* operon was related to the loss of the c3, but it is not the case (**Table S3**).

### A majority of Bcc strains carrying the *hmqABCDEFG* operon also produce HMAQs

We then verified whether the presence of the biosynthetic genes is indicative of known HMAQ production. We cultured the 94 strains that were PCR-positive for *hmqA* and *hmqG* in Tryptic Soy Broth (TSB) medium at 30°C, 250 rpm for overnight under the tested conditions, we could detect the production of HMAQ in 72% of the strains - 65% clinical and 79% environmental - carrying the *hmqABCDEFG* operon (**Figure 3**). None of the PCR-positive *B. anthina, B. diffusa, B. dolosa, B. metallica, B. seminalis* and *B. ubonensis* strains produced HMAQ under our conditions.

**Figure 3.**
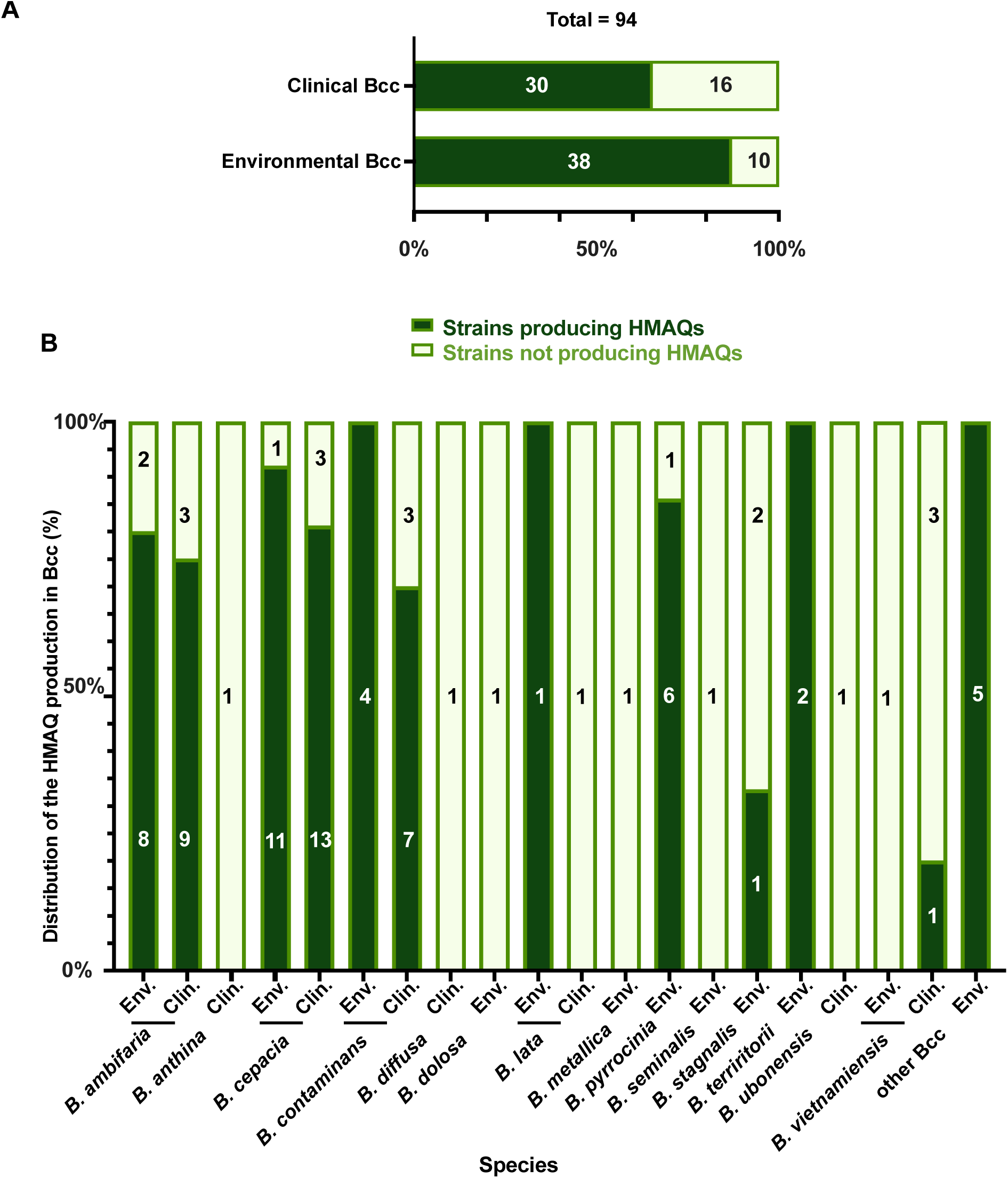
Distribution of the HMAQ production in Bcc. (**A**) The distribution of environmental and clinical Bcc strains regarding their ability to produce HMAQs (B) Distribution of HMAQ producing-Bcc species. HMAQs have been quantified by LC/MS with a limit of detection of 0.05 mg/L for each molecule in the total culture.

To validate our LC-MS method, we measured the production of HMAQs by 31 Bcc strains determined not to carry the *hmqABCDEFG* operon; all the 31 strains – belonging to *B. ambifaria, B. anthina, B. arboris, B. cenocepacia, B. multivorans, B. pyrrocinia, B. stabilis, B. ubonensis* and *B. vietnamiensis* species - did not produce detectable HMAQs (**Table S4**).

HMAQ production was slightly more prevalent among clinical strains for *B. cepacia* and *B. vietnamiensis*. However, it was also more prevalent among environmental strains for *B. ambifaria, B. contaminans*, and *B. lata*. Most of the environmental strains of *B. pyrrocinia, B. stagnalis, B. territorii* and the ‘other Bcc’ group species also produced HMAQ.

### All HMAQ-producing Bcc strains mainly produce the HMAQ-C_7_:2’ and HMAQ-C_9_:2’ congeners

To verify which HMAQ congeners are produced by the various Bcc, we scanned by LC/MS for the 14 congeners of HAQs and HMAQs we had previously identified (**Table S5**) (Vial *et al*., 2008). We found that the major congeners produced were HMAQ-C_7_:2’ and HMAQ-C_9_:2’, as previously observed for *B ambifaria* HSJ1 (Vial *et al*., 2008). Most of the strains were able to produce other HMAQs such as HMAQ-C_7_ and HMAQ-C_8_:2’ (also known as burkholone). HMAQ-C_8_, HMAQ-C_6_ were the most abundant molecules after HMAQ-C_7_:2’ and HMAQ-C_9_:2’ and HHQ-C_9_ was also detected (**Table S6**). Concentrations of HMAQ-C_7_:2’ and HMAQ-C_9_:2’ as well as all identified congeners are listed in **Tables S5 and S6**.

We also found that the concentration of HMAQs produced was variable among the various species. A Kruskal-Wallis test confirmed that there was no statistical difference in the concentrations produced between the clinical and environmental strains (*p*-value of 0.19 for HMAQ-C_7_:2’ and 0.22 for HMAQ-C_9_:2’). However, clinical strains of *B. ambifaria* and *B. vietnamiensis* strains produced more HMAQs than environmental ones. The opposite was observed for *B. cepacia* and *B. contaminans* strains (**Table S6**).

### The presence of the *hmqABCDEFG* and production of HMAQs are not linked to the co-isolation of *P. aeruginosa* nor the origin of samples

Since the *hmqABCDEFG* operon is homologous to the *pqsABCDE* operon in *P. aeruginosa* we wondered if HMAQ production in TSB condition of clinical Bcc was correlated with a co-isolation or a co-localization with *P. aeruginosa* at some point in the patient as well as the origin of the sample (sputum, throat, sinus *etc*.). Information was only available for 53 strains (**Table S7**). Using a Fisher’s Exact Test for Count Data, we did not find a correlation between the presence of the *hmqABCDEFG* operon and the presence of *P. aeruginosa* – at the sampling time or within the previous year – nor with the origin of the sample (**Table 1**).

**Table 1.**
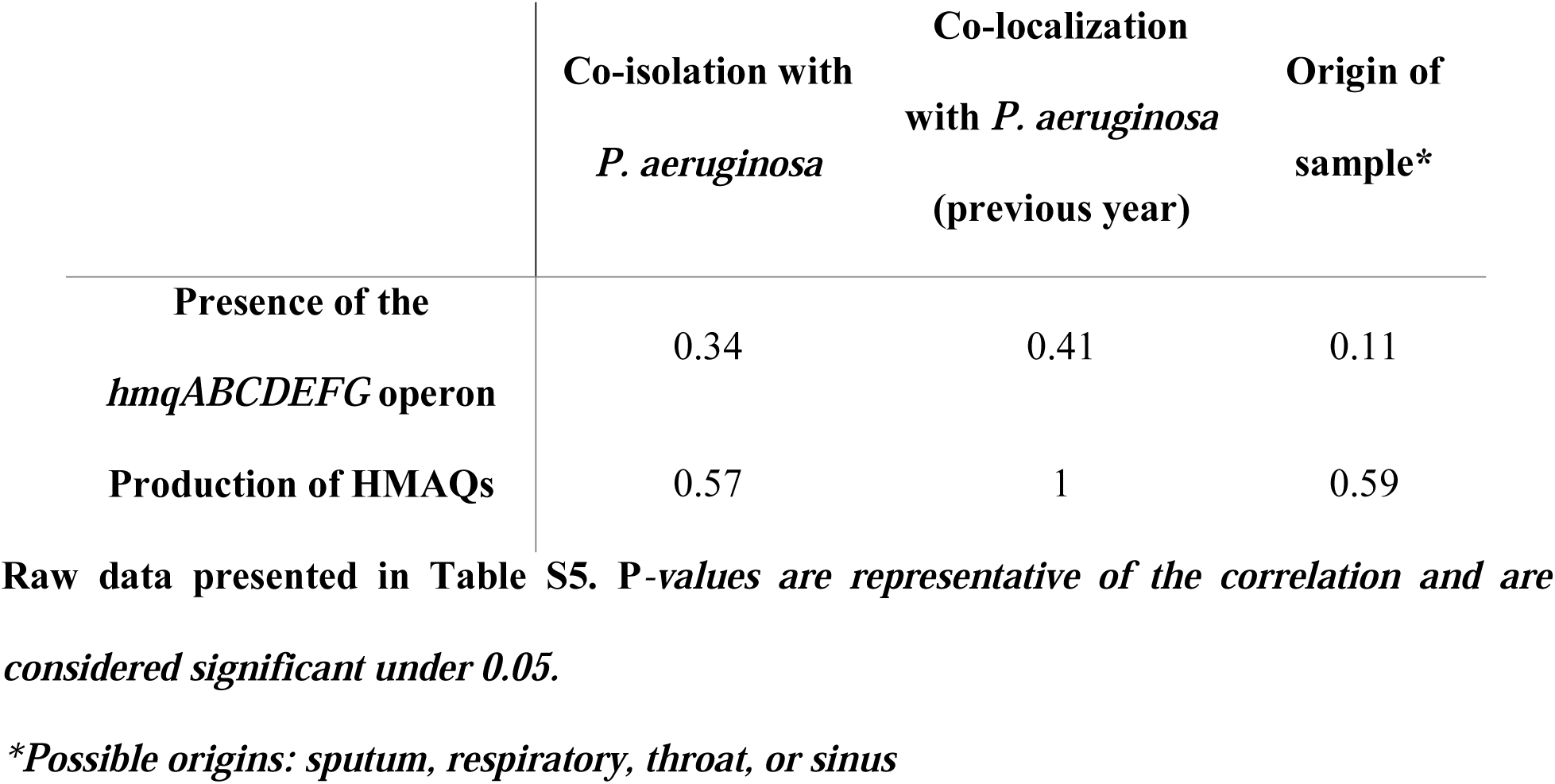
Correlation of the presence of the *hmqABCDEFG* operon and the production of HMAQ in clinical Bcc with sample data

|  | Co-isolation with<br>*P. aeruginosa* | Co-localization<br>with *P. aeruginosa*<br>(previous year) | Origin of<br>sample* |
| --- | --- | --- | --- |
| Presence of the<br>*hmqABCDEFG* operon | 0.34 | 0.41 | 0.11 |
| Production of HMAQs | 0.57 | 1 | 0.59 |

**Raw data presented in Table S5. P-values are representative of the correlation and are considered significant under 0.05.**
**\*Possible origins: sputum, respiratory, throat, or sinus**

### The use of Cystic Fibrosis artificial sputum medium – looking for growth condition relevant for HMAQs production

To induce the production of HMAQs by the 26 Bcc strains carrying the *hmqABCDEFG* operon but for which we could not detect HMAQs when cultured in TSB (at 30°C at 250 rpm, overnight), we assayed the production of these metabolites in Artificial Sputum Medium (ASM; at 30°C at 250 rpm, overnight) and on Tryptic Soy Agar plates (TSA; incubated at 30°C for four days) As shown in **Table 2**, growth in ASM allowed detection of HMAQs in 10 out of the 26 Bcc strains - 7 environmental and 3 clinical strains. Surface growth on TSA plates induced the detectable production of HMAQs for 5 environmental and 4 clinical strains. For the strains already producing HMAQ in TSB most of them also produce HMAQs in ASM and TSA (**Table S8**). These additional culture conditions reduced the number of HMAQ-negative strains to 15 out of 26. Globally, increasing the number of Bcc isolates able to produce HMAQs to a total of 79, that is 84% of strains carrying the *hmqABCDEFG* operon.

**Table 2.** Production of HMAQs in ASM and TSA for strains not producing HMAQs in TSB

| Strains | Type | HMAQs production |  |  |
| --- | --- | --- | --- | --- |
|  |  | TSB | ASM | TSA |
| *B. ambifaria* AMMD | Environmental | - | + | + |
| *B. ambifaria* AU7994 | Clinical | - | - | - |
| *B. ambifaria* CEP1231 | Clinical | - | + | - |
| *B. ambifaria* HI2425 | Environmental | - | + | + |
| *B. ambifaria* VC11631 | Clinical | - | - | - |
| *B. anthina* VC15382 | Clinical | - | - | - |
| *B. cepacia* ATCC25416 | Environmental | - | - | - |
| *B. cepacia* VC13196 | Clinical | - | - | - |
| *B. cepacia* VC13575 | Clinical | - | - | - |
| *B. cepacia* VC19225 | Clinical | - | + | + |
| *B. contaminans* VC15406 | Clinical | - | - | + |
| *B. contaminans* VC16897 | Clinical | - | - | - |
| *B. contaminans* VC16948 | Clinical | - | - | + |
| *B. diffusa* VC14008 | Clinical | - | - | - |
| *B. dolosa* LMG21443 | Environmental | - | - | - |
| *B. lata* VC6377 | Clinical | - | + | + |
| *B. metallica* ES0559 | Environmental | - | - | - |
| *B. pyrrocinia* Bcc indeterminate 9 ES0209 | Environmental | - | + | + |
| *B. seminalis* HI2490 | Environmental | - | + | - |
| *B. stagnalis* Bcc indeterminate 6 HI3537 | Environmental | - | + | + |
| *B. stagnalis* HI2720 | Environmental | - | + | - |
| *B. ubonensis* LMG24263 | Clinical | - | - | - |
| *B. vietnamiensis* CEP0040 | Clinical | - | - | - |
| *B. vietnamiensis* HI3392 | Environmental | - | + | + |
| *B. vietnamiensis* VC17180 | Clinical | - | - | - |
| *B. vietnamiensis* VC9237 | Clinical | - | - | - |

### Low express the *hmqABCDEFG* operon explains absence of HMAQ production in some Bcc strains growing in TSB

Our screening revealed that 26 strains carrying the *hmqABCDEFG* operon do not produce HMAQ in TSB. To investigate the possibility that low transcription of the biosynthetic genes would explain this absence of production, which is compatible with the induction seen when changing the culture conditions, we measured the expression of the *hmqABCDEFG* operon for one HMAQ-negative and one HMAQ-positive strain from each of *B. ambifaria, B. cepacia, B. contaminans* and *B. vietnamiensis* by targeting the *hmqA* gene by RT-PCR.

The results show that HMAQ-negative strains *B. ambifaria* AMMD, *B. cepacia* ATCC25416, *B. contaminans* VC15406, *B. vietnamiensis* VC9237 strains do not express *hmqA* gene when grown in TSB, while *B. ambifaria* HSJ1, *B. cepacia* VC13394, *B. contaminans* FFH2055 and *B. vietnamiensis* VC8245, which produce HMAQs under these conditions, produce a clear *hmqA* transcript (**Table 3; Figure S1**). Extending these results, we hypothesize that the other strains which carry the *hmqABCDEFG* operon and do not produce HMAQs, do not express the *hmqA* gene, at least when grown in TSB at 30°C.

**Table 3.** The expression of the *hmqA* gene in the main species of Bcc strains having the *hmqABCDEFG* operon but which do not produce HMAQs.

| Strains | Type | HMAQs<br>production<br>in TSB | Expression<br>of *hmqA* |
| --- | --- | --- | --- |
| *B. ambifaria* AMMD | Environmental | - | - |
| *B. ambifaria* HSJ1 | Clinical | + | + |
| *B. cepacia* ATCC25416 | Environmental | - | - |
| *B. cepacia* VC13394 | Clinical | + | + |
| *B. contaminans* FFH2055 | Clinical | + | + |
| *B. contaminans* VC15406 | Clinical | - | - |
| *B. vietnamiensis* VC8245 | Clinical | + | + |
| *B. vietnamiensis* VC9237 | Clinical | - | - |

## Discussion

This study aimed at understanding the prevalence of the Hmq system and corresponding HMAQ production in the Bcc to complete or confirm our previous *in silico* analyses (Coulon *et al*., 2019). Only a few strains of *B. ambifaria, B. cepacia, B. pseudomallei* and *B. thailandensis* species were already known to carry the *hmqABCDEFG* operon and to produce HMAQs (Diggle *et al*., 2006; Vial *et al*., 2008; Coulon *et al*., 2019). Even if the role of HMAQs is still unknown, the presence of the *hmqABCDEFG* operon is well-conserved in *B. pseudomallei* and *B. thailandensis* species but remains unclear within the Bcc (Coulon *et al*., 2019). To understand the ecological role of the Hmq system, we first needed to evaluate its distribution. Since available Bcc genomes are not equally distributed between species, we screened 313 Bcc strains for the presence of the *hmqABCDEFG* operon by PCR. We confirmed that the *hmqABCDEFG* operon among the Bcc followed the same distribution when analyzed *in silico* - meaning that a laboratory screening is necessary to complete a bioinformatics study - especially when available whole-genome sequencing is limited. However, we have to take into consideration that our screening could have false-negative results due to the limited availability of whole-genome sequences for some Bcc species (e.g. *B. arboris, B. metallica, B. stabilis*). Nevertheless, our primers were able to amplify *hmqA* and *hmqG* targets in species previously unknown to carry a *hmqABCDEFG* operon (e.g. *B. vietnamiensis*).

Based on our previous results obtained with a few *B. ambifaria* strains, we hypothesized that the Hmq system would be more prevalent among clinical isolates, and produce more HMAQs than environmental ones (Vial *et al*., 2008, 2009; Chapalain *et al*., 2017). Unexpectedly, we uncovered that *B. cenocepacia, B. multivorans* and *B. vietnamiensis* - the prominent Bcc species colonizing immunosuppressed and CF individuals, transmitted between patients (Gilligan, 1991; Speert *et al*., 1994; Govan and Deretic, 1996), do not or rarely (0 out of 72 *B. cenocepacia*, 0 out of 35 *B. multivorans* and 5 out of 37 *B. vietnamiensis*) carry the *hmqABCDEFG* operon. One possibility is that Bcc species more often found as clinical isolates could have lost the Hmq system through evolution and patient-to-patient selection (Coulon *et al*., 2019). Our data suggest that the Hmq system could play a beneficial role in niche adaptation to the rhizosphere microbial community due to the large prevalence of the *hmqABCDEFG* operon among *B. ambifaria, B. cepacia, B. contaminans, B. pyrrocinia* and *B. ubonensis* environmental strains, species known for their preference for the plant root environment (Balandreau *et al*., 2001; Mahenthiralingam *et al*., 2005; Vial et al. 2011; Vidal-Quist *et al*., 2014). The presence of the Hmq system in clinical strains of these five common environmental species is compatible with a recent acquisition in CF population (Huang *et al*., 2001; Mahenthiralingam *et al*., 2005; Loutet and Valvano, 2010). We can also consider the possibility that the presence of competitive microorganisms could favor the production of HMAQs in both clinical and environmental strains - e.g.: *S. aureus* enhances the production of HAQs in *P. aeruginosa* (Michelsen *et al*., 2015).

Since phylogeny does not seem to explain the distribution of the *hmqABCDEFG* operon among Bcc species, we considered the hypothesis that the loss of the c3 replicon, sometimes observed in Bcc phenotypic variants (Agnoli *et al*., 2011), could be related to the absence of the Hmq system, but no correlation was found.

Among the strains having the *hmqABCDEFG* operon, not all produced HMAQs under the tested conditions. It is important to note that the limit of detection of molecules was 0.05 mg/L for each extract which might explain some false-negative results. However, for those strains producing HMAQs in TSB, growth in ASM did not inhibit HMAQs production and stimulated the production in ten additional strains, which could be explained by differences in regulation of the biosynthetic operon. Our results further enforce the need to use nutritionally appropriate media when doing experiments aimed at understanding clinical relevance. For this purpose, we tried to optimize a medium for HMAQs production. However, it was not possible to identify a simple carbon source due to the low production of the molecules in the minimal medium whatever the tested culture time (data not shown).

Lack of transcription of the *hmqABCDEFG* operon seemed to explain that 28% of Bcc strains carrying these genes did not produce HMAQs. Indeed, production could be obtained simply by changing the culture conditions, suggesting that the promotor of the *hmqABCDEFG* was not expressed, although we cannot exclude that it could be nonfunctional in select strains. Future work will involve a better understanding of the nutritional and regulatory elements controlling the expression of the *hmqABCDEFG* operon and production of HMAQs.

Our results suggest that the Hmq system of Bcc is non-essential for pathogenicity but could be required for adaption to a particular environmental niche by, for example, influencing rhizosphere microbial community (Vial *et al*., 2008; Kilani-Feki *et al*., 2011, 2012; Mahenthiralingam *et al*., 2011; Patten *et al*., 2012; Chapalain *et al*., 2013, 2017; Gomes *et al*., 2018; Le Guillouzer, 2018; Jung *et al*., 2018; Mullins *et al*., 2019; Whalen *et al*., 2019; Piochon *et al*., 2020). It will be interesting to investigate a possible correlation between the presence of the *hmqABCDEFG* in Bcc species and their evolutionary trajectory.

## Material and Methods

### Strains and culture conditions

A total of 313 strains isolated from either clinical or environmental settings were used in this study and listed in **Table S1**). Uncertain identification was confirmed by amplifying and sequencing the of the *recA* and *gyrB* genes at the IRCM Sequencing platform (Montreal) following the protocol available on the PubMLST database (https://pubmlst.org/bcc/info/protocol.shtml; **Table S9**).

Strains were cultured in borosilicate tubes containing 3mL tryptic soy broth (TSB) from stocks frozen at -80°C in 15% glycerol and incubated at 30°C with 250 rpm rotative shaking overnight (∼16h).

### Detection of the presence of *hmqABCDEFG* operon by PCR

Genomic DNA was extracted following a previously described method (Durand *et al*., 2015). Briefly, cells were resuspended in lysis buffer (50 mM Tris-HCL pH8, 5 mM EDTA-2Na pH8, 3% SDS) and transferred to a tube containing beads. The cells were lysed in a Fast-Prep-24 instrument (MP Biomedicals, USA). Then the mixes were centrifuged at 8,000 g for 5 min, and the supernatants were transferred to a new tube, and 2.5 N ammonium acetate was added after centrifugation. The supernatant was transferred to a new and one volume isopropanol was added. The pellets were washed by 75% ethanol and dried before to resuspended them into 50 µL water.

For *Burkholderia cepacia* Research Laboratory and Repository and Canadian *Burkholderia cepacia* complex Research and Referral Repository strains, genomic DNA was extracted using a 96-wells plate gDNA extraction kit (Favorgen, Canada).

The *hmqA* and *hmqG* genes were amplified by PCR using the EasyTaq polymerase (Transgen, Canada). Primers were designed based on a consensus sequence of 11 complete Bcc sequences (**Table S9**). A strain was considered to have a complete *hmqABCDEFG* operon when amplification for both *hmqA* and *hmqG* were obtained.

### Quantification of HMAQ production by LC/MS/MS

Bcc strains were cultured in 5 mL tryptic soy broth (TSB) at a starting OD_600nm_ of 0.05 and incubated at 30°C with shaking for an overnight. 5,6,7,8-tetradeutero-4-hydroxy-2- heptylquinoline (HHQ-d4, Sigma) was used as an internal standard. The total HMAQ were extracted from 4 mL culture with one volume of ethyl acetate. After nitrogen evaporation, the extracts were resuspended in 400 μL of HPLC-grade acetonitrile. Samples were analyzed by liquid chromatography coupled with a mass spectrometer (LC/MS) in positive electrospray ionization using a Kinetex 5 μM EVO C18 100 Å 100×3 mm reverse phase column as previously described by (Vial *et al*., 2008). A Quattro Premier XE triple quadrupole was used as a detector (Waters). A full scan mode with a scanning range of 130 to 350 Da and a multiple reaction monitoring (MRM) program were used to detected HMAQ families based on (Vial *et al*., 2008). This experiment was conducted with three independent biological replicates.

### Artificial Sputum medium (ASM) and Tryptic soy Agar (TSA) medium assay

The strains were grown in ASM and TSA plate out from overnight cultures and incubated at 30°C for 24h and four days, respectively.

One mL of ASM culture were extracted as previously described in HMAQ production method. For each TSA plate, 5 mL water was added to be able to extract HMAQs from the agar. For each sample, 1 mL was extracted by one volume ethyl acetate containing 4 ppm HHQ-D4, concentrated 10 times, resuspended in HPLC-grade acetonitrile and analyzed as previously described in HMAQ extraction method.

The experiments were performed in two independent biological replicates.

### Detection of the expression of the *hmqABCDEFG* operon by RT-PCR

Total RNA was extracted from cultures grown in TSB to an OD_600_ of 3.0 using Transzol (Transgene, Canada) by following the manufacturer’s instruction. Residual DNA was removed using the Turbo DNAse (Thermo Fisher, Canada). Reverse-transcription was performed using the I-Script kit (BioRad, Canada). The expression of the *hmqABCDEFG* operon gene was determined by PCR targeting the *hmqA* and *ndh* as a reference gene (**Table S9)** (Subsin *et al*., 2007).

To determine the *hmqA* gene primers, a semi-random PCR was performed on genomic DNA from *B. cepacia* VC13394, *B. contaminans* VC15406, *B. vietnamiensis* VC8245 and *B. vietnamiensis* VC9237 using the two specific primers (*hmqA*_semirandom_3F and *hmqA*_semirandom_2F) and sequencing of the PCR product using a third primer (*hmqA*_semirandom_F; **Table S3**) by following the protocol of Jacobs *et al*. (2003).

### The link of the presence of the *hmqABCDEFG* operon and the production of HMAQ in Bcc with different characteristics

Based on our qualitative data (**Tables 1 and S7**), we studied the correlation by Fisher’s exact test for count data using R software (http://www.R-project.org ; (Team, 2018)).

## Acknowledgments

We thank Dr Silvia Cardona for providing *B. cepacia* BTS13 strain; Dr David Wagner for *B. stagnalis* MSMB1956WGS, *B. territorii* MSMB1301WGS and *B. territorii* MSMB1502WGS and Dr John LiPuma for providing the 75 environmental strains listed in **Table S1**.

We also want to thank Pr Phillipe Constant for helping with statistics, and Marie-Christine Groleau for answering to experiments wonders and for her great work on reviewing the article draft.

