## Supplemental data for "Presence of the Hmq system and production of 4-hydroxy-3-methyl-2-alkylquinolines is heterogeneously distributed between *Burkholderia cepacia* complex species and more prevalent among environmental than clinical isolates"

**Table S1. List of strains used in this study.**

| Strains | Type | References | Other names |
|---|---|---|---|
| ***Burkholderia ambifaria*** | | | |
| *B. ambifaria* AMMD | Environmental isolate | Coenye et al. 2001 | LMG 19182/FC0768/BCC0588 |
| *B. ambifaria* AU0212 | CF isolate (USA) | Payne et al. 2005 | |
| *B. ambifaria* AU4157 | Clinical isolate | BcRLR | |
| *B. ambifaria* AU7994 | Clinical isolate | | |
| *B. ambifaria* AU8235 | Clinical isolate | BcRLR | |
| *B. ambifaria* CEP0516 | CF isolate (Australia) | Coenye et al. 2001 | |
| *B. ambifaria* CEP0617 | Clinical isolate | Coenye et al. 2001 | LMG-P 24636 |
| *B. ambifaria* CEP0958 | CF isolate (Australia) | Coenye et al. 2001 | |
| *B. ambifaria* CEP0990 | Clinical isolate | | |
| *B. ambifaria* CEP0996 | CF isolate (Australia) | Coenye et al. 2003 | LMG 19467 |
| *B. ambifaria* CEP1231 | Clinical isolate | | |
| *B. ambifaria* ES0020 | Environmental isolate (USA) | BcRLR | |
| *B. ambifaria* HI2425 | Environmental isolate (USA) | BcRLR | |
| *B. ambifaria* HI2468 | Environmental isolate (USA) | BcRLR | |
| *B. ambifaria* HI2482 | Environmental isolate (USA) | BcRLR | |
| *B. ambifaria* HI2626 | Environmental isolate (USA) | BcRLR | |
| *B. ambifaria* HI2672 | Environmental isolate (USA) | BcRLR | |
| *B. ambifaria* HI3544 | Environmental isolate (USA) | BcRLR | |
| *B. ambifaria* HI3590 | Environmental isolate (USA) | BcRLR | |
| *B. ambifaria* HI3687 | Environmental isolate (USA) | BcRLR | |
| *B. ambifaria* HI3709 | Environmental isolate (USA) | BcRLR | |
| *B. ambifaria* HI3738 | Environmental isolate (USA) | BcRLR | |
| *B. ambifaria* HI3890 | Environmental isolate (USA) | BcRLR | |
| *B. ambifaria* HSJ1 | CF isolate isolate (Canada) | Vial et al. 2008 | |
| *B. ambifaria* IOP40-10 | Environmental isolate | J. Tiedje collection | |
| *B. ambifaria* LMG17828 | | Coenye et al. 2001 | ATCC 53266/FC0662 |
| *B. ambifaria* MW2073 | Environmental isolate | | |
| *B. ambifaria* PC736 | Environmental isolate (USA) | BcRLR | |
| *B. ambifaria* PHP7 | Environmental isolate | Coenye et al. 2001 | |
| *B. ambifaria* VC11631 | CF isolate (Canada) | CBCCRRR | |
| *B. ambifaria* VC15422 | CF isolate (Canada) | CBCCRRR | |
| *B. ambifaria* VC16196 | Clinical isolate (Canada) | CBCCRRR | |
| ***Burkholderia anthina*** | | | |
| *B. anthina* Bcc indeterminate 7 HI3538 | Environmental isolate (USA) | BcRLR | |
| *B. anthina* HI3655 | Environmental isolate (USA) | BcRLR | |
| *B. anthina* VC15382 | Clinical isolate (Canada) | CBCCRRR | |
| *B. anthina* VC16083 | Clinical isolate (Canada) | CBCCRRR | |
| ***Burkholderia arboris*** | | | |
| *B. arboris* ES0222 | Environmental isolate (USA) | BcRLR | |
| *B. arboris* ES0263 | Environmental isolate (USA) | BcRLR | |
| *B. arboris* VC10224 | Clinical isolate (Canada) | CBCCRRR | |
| *B. arboris* VC8833 | CF isolate isolate (Canada) | CBCCRRR | |
| ***Burkholderia cenocepacia*** | | | |
| *B. cenocepacia* CEP024 | CF isolate isolate (Canada) | Speert collection | |
| *B. cenocepacia* CEP0511 | CF isolate isolate (Australia) | Baldwin et al. 2005 | LMG 18830 |
| *B. cenocepacia* CEP0565 | Clinical isolate | | |
| *B. cenocepacia* ES1405 | Environmental isolate (USA) | BcRLR | |
| *B. cenocepacia* HI2424 | Environmental isolate (USA) | BcRLR | |
| *B. cenocepacia* HI2606 | Environmental isolate (USA) | BcRLR | |
| *B. cenocepacia* HI2876 | Environmental isolate (USA) | BcRLR | |
| *B. cenocepacia* HI2976 | Environmental isolate (USA) | BcRLR | |
| *B. cenocepacia* HI3540 | Environmental isolate (USA) | BcRLR | |
| *B. cenocepacia* HI3855 | Environmental isolate (USA) | BcRLR | |
| *B. cenocepacia* HI4004 | Environmental isolate (USA) | BcRLR | |
| *B. cenocepacia* HI4101 | Environmental isolate (USA) | BcRLR | |
| *B. cenocepacia* HI4143 | Environmental isolate (USA) | BcRLR | |
| *B. cenocepacia* HI4261 | Environmental isolate (USA) | BcRLR | |
| *B. cenocepacia* HI4437 | Environmental isolate (USA) | BcRLR | |
| *B. cenocepacia* HI4904 | Environmental isolate (USA) | BcRLR | |
| *B. cenocepacia* K56-2 | CF isolate isolate (Canada) | LiPuma et al. 2001 | |
| *B. cenocepacia* IIIA VC10277 | CF isolate (Canada) | CBCCRRR | |
| *B. cenocepacia* IIIA VC12308 | CF isolate (Canada) | CBCCRRR | |
| *B. cenocepacia* IIIA VC13139 | CF isolate (Canada) | CBCCRRR | |
| *B. cenocepacia* IIIA VC14610 | CF isolate (Canada) | CBCCRRR | |
| *B. cenocepacia* IIIA VC15419 | CF isolate (Canada) | CBCCRRR | |
| *B. cenocepacia* IIIA VC15451 | CF isolate (Canada) | CBCCRRR | |
| *B. cenocepacia* IIIA VC16156 | CF isolate (Canada) | CBCCRRR | |
| *B. cenocepacia* IIIA VC16199 | CF isolate (Canada) | CBCCRRR | |
| *B. cenocepacia* IIIA VC16873 | CF isolate (Canada) | CBCCRRR | |
| *B. cenocepacia* IIIA VC16874 | CF isolate (Canada) | CBCCRRR | |
| *B. cenocepacia* IIIA VC17671 | CF isolate (Canada) | CBCCRRR | |
| *B. cenocepacia* IIIA VC17819 | CF isolate (Canada) | CBCCRRR | |
| *B. cenocepacia* IIIA VC18585 | CF isolate (Canada) | CBCCRRR | |
| *B. cenocepacia* IIIA VC18609 | CF isolate (Canada) | CBCCRRR | |
| *B. cenocepacia* IIIA VC18996 | CF isolate (Canada) | CBCCRRR | |
| *B. cenocepacia* IIIA VC18999 | CF isolate (Canada) | CBCCRRR | |
| *B. cenocepacia* IIIA VC3917 | CF isolate (Canada) | CBCCRRR | |
| *B. cenocepacia* IIIA VC5069 | CF isolate (Canada) | CBCCRRR | |
| *B. cenocepacia* IIIA VC5621 | CF isolate (Canada) | CBCCRRR | |
| *B. cenocepacia* IIIA VC6356 | CF isolate (Canada) | CBCCRRR | |
| *B. cenocepacia* IIIA VC6553 | CF isolate (Canada) | CBCCRRR | |
| *B. cenocepacia* IIIA VC8286 | CF isolate (Canada) | CBCCRRR | |
| *B. cenocepacia* IIIA VC8356 | CF isolate (Canada) | CBCCRRR | |
| *B. cenocepacia* IIIA VC8426 | CF isolate (Canada) | CBCCRRR | |
| *B. cenocepacia* IIIA VC8607 | CF isolate (Canada) | CBCCRRR | |
| *B. cenocepacia* IIIA VC8611 | CF isolate (Canada) | CBCCRRR | |
| *B. cenocepacia* IIIA VC8614 | CF isolate (Canada) | CBCCRRR | |
| *B. cenocepacia* IIIA VC9080 | CF isolate (Canada) | CBCCRRR | |
| *B. cenocepacia* IIIA VC9296 | CF isolate (Canada) | CBCCRRR | |
| *B. cenocepacia* IIIB VC11311 | CF isolate (Canada) | CBCCRRR | |
| *B. cenocepacia* IIIB VC11653 | CF isolate (Canada) | CBCCRRR | |

| | | | |
|---|---|---|---|
| *B. cenocepacia* IIIB VC13104 | CF isolate (Canada) | CBCCRRR | |
| *B. cenocepacia* IIIB VC13187 | CF isolate (Canada) | CBCCRRR | |
| *B. cenocepacia* IIIB VC14376 | CF isolate (Canada) | CBCCRRR | |
| *B. cenocepacia* IIIB VC14524 | CF isolate (Canada) | CBCCRRR | |
| *B. cenocepacia* IIIB VC14529 | CF isolate (Canada) | CBCCRRR | |
| *B. cenocepacia* IIIB VC15122 | CF isolate (Canada) | CBCCRRR | |
| *B. cenocepacia* IIIB VC15240 | CF isolate (Canada) | CBCCRRR | |
| *B. cenocepacia* IIIB VC15241 | CF isolate (Canada) | CBCCRRR | |
| *B. cenocepacia* IIIB VC16932 | CF isolate (Canada) | CBCCRRR | |
| *B. cenocepacia* IIIB VC17657 | CF isolate (Canada) | CBCCRRR | |
| *B. cenocepacia* IIIB VC18097 | CF isolate (Canada) | CBCCRRR | |
| *B. cenocepacia* IIIB VC18107 | CF isolate (Canada) | CBCCRRR | |
| *B. cenocepacia* IIIB VC18236 | CF isolate (Canada) | CBCCRRR | |
| *B. cenocepacia* IIIB VC18454 | CF isolate (Canada) | CBCCRRR | |
| *B. cenocepacia* IIIB VC18569 | CF isolate (Canada) | CBCCRRR | |
| *B. cenocepacia* IIIB VC18658 | CF isolate (Canada) | CBCCRRR | |
| *B. cenocepacia* IIIB VC5625 | CF isolate (Canada) | CBCCRRR | |
| *B. cenocepacia* IIIB VC6598 | CF isolate (Canada) | CBCCRRR | |
| *B. cenocepacia* IIIB VC7349 | CF isolate (Canada) | CBCCRRR | |
| *B. cenocepacia* IIIB VC7849 | CF isolate (Canada) | CBCCRRR | |
| *B. cenocepacia* IIIB VC7911 | CF isolate (Canada) | CBCCRRR | |
| *B. cenocepacia* IIIB VC8340 | CF isolate (Canada) | CBCCRRR | |
| *B. cenocepacia* IIIB VC8870 | CF isolate (Canada) | CBCCRRR | |
| *B. cenocepacia* IIIB VC9859 | CGD (Canada) | CBCCRRR | |
| ***Burkholderia cepacia*** | | | |
| *B. cepacia* ATCC25416 | Environmental isolate | Yabuuchi et al. 1992 | LMG1222/CEP0031 |
| *B. cepacia* CEP0509 | CF isolate (Australia) | Vandamme et al. 1997 | LMG18821 |
| *B. cepacia* BTS13 | CF isolate (Italy) | Lagatolla et al. 2002 | |
| *B. cepacia* HI2430 | Environmental isolate (USA) | BcRLR | |
| *B. cepacia* HI2563 | Environmental isolate (USA) | BcRLR | |
| *B. cepacia* HI2578 | Environmental isolate (USA) | BcRLR | |
| *B. cepacia* HI2615 | Environmental isolate (USA) | BcRLR | |
| *B. cepacia* HI2671 | Environmental isolate (USA) | BcRLR | |
| *B. cepacia* HI2741 | Environmental isolate (USA) | BcRLR | |
| *B. cepacia* HI3312 | Environmental isolate (USA) | BcRLR | |
| *B. cepacia* HI3551 | Environmental isolate (USA) | BcRLR | |
| *B. cepacia* HI3708 | Environmental isolate (USA) | BcRLR | |
| *B. cepacia* HI3895 | Environmental isolate (USA) | BcRLR | |
| *B. cepacia* HI4577 | Environmental isolate (USA) | BcRLR | |
| *B. cepacia* VC13132 | CF isolate (Canada) | CBCCRRR | |
| *B. cepacia* VC13196 | CF isolate (Canada) | CBCCRRR | |
| *B. cepacia* VC13394 | CF isolate (Canada) | CBCCRRR | |
| *B. cepacia* VC13575 | CF isolate (Canada) | CBCCRRR | |
| *B. cepacia* VC14106 | CF isolate (Canada) | CBCCRRR | |
| *B. cepacia* VC14457 | CF isolate (Canada) | CBCCRRR | |
| *B. cepacia* VC16383 | Clinical isolate (Canada) | CBCCRRR | |
| *B. cepacia* VC16708 | CF isolate (Canada) | CBCCRRR | |
| *B. cepacia* VC17333 | CF isolate (Canada) | CBCCRRR | |
| *B. cepacia* VC17746 | NON CF isolate (Canada) | CBCCRRR | |
| *B. cepacia* VC17928 | CF isolate (Canada) | CBCCRRR | |
| *B. cepacia* VC18315 | CF isolate isolate (Canada) | CBCCRRR | |
| *B. cepacia* VC18839 | NON CF isolate (Canada) | CBCCRRR | |
| *B. cepacia* VC18842 | NON CF isolate (Canada) | CBCCRRR | |
| *B. cepacia* VC19225 | CF isolate (Canada) | CBCCRRR | |
| *B. cepacia* VC19276 | CF isolate (Canada) | CBCCRRR | |
| *B. cepacia* VC9490 | CF isolate (Canada) | CBCCRRR | |
| ***Burkholderia contaminans*** | | | |
| *B. contaminans* FFH2055 | CF isolate (Argentina) | Nunvar et al. 2016 | |
| *B. contaminans* HI3422 | Environmental isolate (USA) | BcRLR | |
| *B. contaminans* HI3570 | Environmental isolate (USA) | BcRLR | |
| *B. contaminans* HI3852 | Environmental isolate (USA) | BcRLR | |
| *B. contaminans* HI3887 | Environmental isolate (USA) | BcRLR | |
| *B. contaminans* HI4067 | Environmental isolate (USA) | BcRLR | |
| *B. contaminans* HI4232 | Environmental isolate (USA) | BcRLR | |
| *B. contaminans* HI4402 | Environmental isolate (USA) | BcRLR | |
| *B. contaminans* VC14347 | CF isolate (Canada) | CBCCRRR | |
| *B. contaminans* VC15406 | CF isolate (Canada) | CBCCRRR | |
| *B. contaminans* VC16087 | CF isolate (Canada) | CBCCRRR | |
| *B. contaminans* VC16848-b | CF isolate (Canada) | CBCCRRR | |
| *B. contaminans* VC16897 | (Canada) | CBCCRRR | |
| *B. contaminans* VC16948 | CF isolate (Canada) | CBCCRRR | |
| *B. contaminans* VC19056 | CF isolate (Canada) | CBCCRRR | |
| *B. contaminans* VC19124 | CF isolate (Canada) | CBCCRRR | |
| *B. contaminans* VC9624 | CF isolate (Canada) | CBCCRRR | |
| ***Burkholderia diffusa*** | | | |
| *B. diffusa* HI2617 | Environmental isolate (USA) | BcRLR | |
| *B. diffusa* HI3576 | Environmental isolate (USA) | BcRLR | |
| *B. diffusa* HI3672 | Environmental isolate (USA) | BcRLR | |
| *B. diffusa* HI3740 | Environmental isolate (USA) | BcRLR | |
| *B. diffusa* VC14008 | CF isolate (Canada) | CBCCRRR | |
| *B. diffusa* VC15063 | CF isolate (Canada) | CBCCRRR | |
| *B. diffusa* VC6752 | CF isolate (Canada) | CBCCRRR | |
| *B. diffusa* VC6966 | NON CF isolate (Canada) | CBCCRRR | |
| *B. diffusa* VC7394 | CF isolate (Canada) | CBCCRRR | |
| *B. diffusa* VC7913 | CF isolate (Canada) | CBCCRRR | |
| ***Burkholderia dolosa*** | | | |
| *B. dolosa* CEP0021 | CF isolate (Canada) | CBCCRRR | |
| *B. dolosa* LMG21443 | Environmental isolate | Vandamme et al. 2002 | |
| *B. dolosa* VC14902 | CF isolate (Canada) | CBCCRRR | |
| *B. dolosa* VC17647 | CF isolate (Canada) | CBCCRRR | |
| ***Burkholderia lata*** | | | |
| *B. lata* BC01 | Environmental isolate (USA) | BcRLR | |
| *B. lata* VC19230 | (Canada) | CBCCRRR | |
| *B. lata* VC6377 | CF isolate (Canada) | CBCCRRR | |

| | | | |
|---|---|---|---|
| *B. lata* VC8171 | CF isolate (Canada) | CBCCRRR | |
| ***Burkholderia metallica*** | | | |
| *B. metallica* ES0559 | Environmental isolate (USA) | BcRLR | |
| *B. metallica* HI3647 | Environmental isolate (USA) | BcRLR | |
| *B. metallica* VC15467 | CF isolate (Canada) | CBCCRRR | |
| *B. metallica* VC8135 | CF isolate (Canada) | CBCCRRR | |
| ***Burkholderia multivorans*** | | | |
| *B. multivorans* LMG16660 | CF isolate isolate (Canada) | CBCCRRR | CEP0781 |
| *B. multivorans* HI2790 | Environmental isolate (USA) | BcRLR | |
| *B. multivorans* LMG17588 | Environmental isolate | Vandamme et al. 1997 | ATCC17616/CEP0144 |
| *B. multivorans* VC12152 | CF isolate (Canada) | CBCCRRR | |
| *B. multivorans* VC12258 | CF isolate (Canada) | CBCCRRR | |
| *B. multivorans* VC12539 | CF isolate (Canada) | CBCCRRR | |
| *B. multivorans* VC12675 | Clinical isolate (Canada) | CBCCRRR | |
| *B. multivorans* VC13125 | CF isolate (Canada) | CBCCRRR | |
| *B. multivorans* VC13145 | CF isolate (Canada) | CBCCRRR | |
| *B. multivorans* VC13162 | CF isolate (Canada) | CBCCRRR | |
| *B. multivorans* VC13451 | NON CF isolate (Canada) | CBCCRRR | |
| *B. multivorans* VC13673 | CF isolate (Canada) | CBCCRRR | |
| *B. multivorans* VC13702 | CF isolate (Canada) | CBCCRRR | |
| *B. multivorans* VC13776 | CF isolate (Canada) | CBCCRRR | |
| *B. multivorans* VC14090 | CF isolate (Canada) | CBCCRRR | |
| *B. multivorans* VC14422 | NON CF isolate (Canada) | CBCCRRR | |
| *B. multivorans* VC14443 | CF isolate (Canada) | CBCCRRR | |
| *B. multivorans* VC14749 | CF isolate (Canada) | CBCCRRR | |
| *B. multivorans* VC14757 | CF isolate (Canada) | CBCCRRR | |
| *B. multivorans* VC15002 | CF isolate (Canada) | CBCCRRR | |
| *B. multivorans* VC15085 | CF isolate (Canada) | CBCCRRR | |
| *B. multivorans* VC15268 | CF isolate (Canada) | CBCCRRR | |
| *B. multivorans* VC15273 | CF isolate (Canada) | CBCCRRR | |
| *B. multivorans* VC15814 | CF isolate (Canada) | CBCCRRR | |
| *B. multivorans* VC15834 | CF isolate (Canada) | CBCCRRR | |
| *B. multivorans* VC15873 | CF isolate (Canada) | CBCCRRR | |
| *B. multivorans* VC15952 | CF isolate (Canada) | CBCCRRR | |
| *B. multivorans* VC15953 | CF isolate (Canada) | CBCCRRR | |
| *B. multivorans* VC15977 | CF isolate (Canada) | CBCCRRR | |
| *B. multivorans* VC16475 | CF isolate (Canada) | CBCCRRR | |
| *B. multivorans* VC16487 | CF isolate (Canada) | CBCCRRR | |
| *B. multivorans* VC16759 | CF isolate (Canada) | CBCCRRR | |
| *B. multivorans* VC16959 | CF isolate (Canada) | CBCCRRR | |
| *B. multivorans* VC17546 | Clinical isolate (Canada) | CBCCRRR | |
| *B. multivorans* VC18625 | CF isolate (Canada) | CBCCRRR | |
| *B. multivorans* VC3419 | CF isolate (Canada) | CBCCRRR | |
| *B. multivorans* VC4282 | CF isolate (Canada) | CBCCRRR | |
| *B. multivorans* VC6534 | CF isolate (Canada) | CBCCRRR | |
| *B. multivorans* VC6564 | CF isolate (Canada) | CBCCRRR | |
| *B. multivorans* VC7102 | CF isolate (Canada) | CBCCRRR | |
| *B. multivorans* VC7704 | CF isolate (Canada) | CBCCRRR | |
| *B. multivorans* VC7870 | CF isolate (Canada) | CBCCRRR | |
| *B. multivorans* VC7960 | CF isolate (Canada) | CBCCRRR | |
| *B. multivorans* VC9159 | CF isolate (Canada) | BcRLR | |
| *B. multivorans* VC9858 | CF isolate (Canada) | CBCCRRR | |
| ***Burkholderia pyrrocinia*** | | | |
| *B. pyrrocinia* CH-67 | Environmental isolate (Korea) | Lee et al. 2011 | |
| *B. pyrrocinia* Bcc indeterminate 1 ES0490 | Environmental isolate (USA) | BcRLR | |
| *B. pyrrocinia* Bcc indeterminate 2 BC02 | Environmental isolate (USA) | BcRLR | |
| *B. pyrrocinia* Bcc indeterminate 5 HI2575 | Environmental isolate (USA) | BcRLR | |
| *B. pyrrocinia* Bcc indeterminate 5 HI2690 | Environmental isolate (USA) | BcRLR | |
| *B. pyrrocinia* Bcc indeterminate 5 HI2701 | Environmental isolate (USA) | BcRLR | |
| *B. pyrrocinia* Bcc indeterminate 5 HI3892 | Environmental isolate (USA) | BcRLR | |
| *B. pyrrocinia* Bcc indeterminate 9 ES0209 | Environmental isolate (USA) | BcRLR | |
| *B. pyrrocinia* LMG21824 | CF isolate (Canada) | Coenye et al. 2003 | |
| ***Burkholderia seminalis*** | | | |
| *B. seminalis* HI2490 | Environmental isolate (USA) | BcRLR | |
| ***Burkholderia stabilis*** | | | |
| *B. stabilis* C7322 | CF isolate (Canada) | Mahenthiralingam et al. 2000 | |
| *B. stabilis* HI2462 | Environmental isolate (USA) | BcRLR | |
| *B. stabilis* VC10097 | NON CF isolate (Canada) | CBCCRRR | |
| *B. stabilis* VC12965 | Environmental isolate (Canada) | CBCCRRR | |
| *B. stabilis* VC12344 | CF isolate (Canada) | CBCCRRR | |
| *B. stabilis* VC17755 | Cancer (Canada) | CBCCRRR | |
| *B. stabilis* VC6296 | CF isolate (Canada) | CBCCRRR | |
| *B. stabilis* VC6482 | CF isolate (Canada) | CBCCRRR | |
| *B. stabilis* VC6747 | CF isolate (Canada) | CBCCRRR | |
| *B. stabilis* VC6749 | CF isolate (Canada) | CBCCRRR | |
| *B. stabilis* VC6753 | CF isolate (Canada) | CBCCRRR | |
| *B. stabilis* VC8622 | CF isolate (Canada) | CBCCRRR | |
| *B. stabilis* VC8623 | CF isolate (Canada) | CBCCRRR | |
| *B. stabilis* VC8629 | CF isolate (Canada) | CBCCRRR | |
| *B. stabilis* VC8636 | CF isolate (Canada) | CBCCRRR | |
| *B. stabilis* VC8638 | CF isolate (Canada) | CBCCRRR | |
| *B. stabilis* VC8967 | NON CF isolate (Canada) | CBCCRRR | |
| *B. stabilis* VC8971 | CF isolate (Canada) | CBCCRRR | |
| *B. stabilis* VC9042 | CF isolate (Canada) | CBCCRRR | |
| *B. stabilis* VC9562 | CF isolate (Canada) | CBCCRRR | |
| *B. stabilis* VC9945 | CF isolate (Canada) | CBCCRRR | |
| ***Burkholderia stagnalis*** | | | |
| *B. stagnalis* HI2720 | Environmental isolate (USA) | BcRLR | |
| *B. stagnalis* Bcc indeterminate 6 HI3537 | Environmental isolate (USA) | BcRLR | |
| *B. stagnalis* MSMB1956WGS | Environmental isolate (USA) | BDEC | |
| ***Burkholderia territorii*** | | | |
| *B. territorii* MSMB1301WGS | Environmental isolate (USA) | BDEC | |
| *B. territorii* MSMB1502WGS | Environmental isolate (USA) | BDEC | |
| ***Burkholderia ubonensis*** | | | |

| | | | |
|---|---|---|---|
| *B. ubonensis* LMG20358 | Environmental isolate (Thailand) | Coenye et al. 2001 | BCC1603 |
| *B. ubonensis* LMG24263 | Nosocomial (Thailand) | Vanlaere et al. 2008 | |
| ***Burkholderia vietnamiensis*** | | | |
| *B. vietnamiensis* CEP0040 | CF isolate isolate (Canada) | Mahenthiraligham collection | LMG 18835 |
| *B. vietnamiensis* G4 | Environmental isolate | Nelson et al. 1987 | |
| *B. vietnamiensis* HI3392 | Environmental isolate (USA) | BcRLR | |
| *B. vietnamiensis* HI3534 | Environmental isolate (USA) | BcRLR | |
| *B. vietnamiensis* VC0024 | CF isolate isolate (Canada) | CBCCRRR | |
| *B. vietnamiensis* VC10362 | CF isolate isolate (Canada) | CBCCRRR | |
| *B. vietnamiensis* VC10442 | NON CF isolate isolate (Canada) | CBCCRRR | |
| *B. vietnamiensis* VC10676 | CF isolate (Canada) | CBCCRRR | |
| *B. vietnamiensis* VC11253 | CF isolate (Canada) | CBCCRRR | |
| *B. vietnamiensis* VC11275 | CF isolate (Canada) | CBCCRRR | |
| *B. vietnamiensis* VC11668 | CF isolate (Canada) | CBCCRRR | |
| *B. vietnamiensis* VC12002 | CF isolate (Canada) | CBCCRRR | |
| *B. vietnamiensis* VC13138 | CF isolate (Canada) | CBCCRRR | |
| *B. vietnamiensis* VC13308 | CF isolate (Canada) | CBCCRRR | |
| *B. vietnamiensis* VC13830 | CF isolate (Canada) | CBCCRRR | |
| *B. vietnamiensis* VC13984 | CF isolate (Canada) | CBCCRRR | |
| *B. vietnamiensis* VC14091 | CF isolate (Canada) | CBCCRRR | |
| *B. vietnamiensis* VC14473 | CF isolate (Canada) | CBCCRRR | |
| *B. vietnamiensis* VC14737 | CF isolate (Canada) | CBCCRRR | |
| *B. vietnamiensis* VC15208 | CF isolate (Canada) | CBCCRRR | |
| *B. vietnamiensis* VC15292 | CF isolate (Canada) | CBCCRRR | |
| *B. vietnamiensis* VC15774 | CF isolate (Canada) | CBCCRRR | |
| *B. vietnamiensis* VC16431 | CF isolate (Canada) | CBCCRRR | |
| *B. vietnamiensis* VC17180 | CF isolate (Canada) | CBCCRRR | |
| *B. vietnamiensis* VC17270 | CF isolate (Canada) | CBCCRRR | |
| *B. vietnamiensis* VC17399 | CF isolate (Canada) | CBCCRRR | |
| *B. vietnamiensis* VC17834 | CF isolate (Canada) | CBCCRRR | |
| *B. vietnamiensis* VC18210 | CF isolate (Canada) | CBCCRRR | |
| *B. vietnamiensis* VC18530 | CF isolate (Canada) | CBCCRRR | |
| *B. vietnamiensis* VC18712 | CF isolate (Canada) | CBCCRRR | |
| *B. vietnamiensis* VC18844 | CF isolate (Canada) | CBCCRRR | |
| *B. vietnamiensis* VC2824 | CF isolate (Canada) | CBCCRRR | |
| *B. vietnamiensis* VC5914 | CF isolate (Canada) | CBCCRRR | |
| *B. vietnamiensis* VC8245 | CF isolate (Canada) | CBCCRRR | |
| *B. vietnamiensis* VC8613 | CF isolate (Canada) | CBCCRRR | |
| *B. vietnamiensis* VC9237 | CF isolate (Canada) | CBCCRRR | |
| *B. vietnamiensis* VC9752 | CF isolate (Canada) | CBCCRRR | |
| **Other Bcc group** | | | |
| *B.* sp LMI-SB2 | | | |
| *B.* spp VC14128 | CF isolate (Canada) | CBCCRRR | |
| *B.* spp VC15804 | CF isolate (Canada) | CBCCRRR | |
| *B.* spp VC16512 | (Canada) | CBCCRRR | |
| *B.* spp VC18848 | CF isolate (Canada) | CBCCRRR | |
| Other Bcc - Bcc indeterminate 1 BC06 | Environmental isolate (USA) | BcRLR | |
| Other Bcc - Bcc indeterminate 3 BC13 | Environmental isolate (USA) | BcRLR | |
| Other Bcc - Bcc indeterminate 3 ES0139 | Environmental isolate (USA) | BcRLR | |
| Other Bcc - Bcc indeterminate 4 BC04 | Environmental isolate (USA) | BcRLR | |
| Other Bcc - Bcc indeterminate 5 BC03 | Environmental isolate (USA) | BcRLR | |
| Other Bcc - Bcc indeterminate 8 HI4407 | Environmental isolate (USA) | BcRLR | |

CBCCRRR: Strains were provided by Canadian *Burkholderia cepacia complex* Research and Referral Repository, University of British Colombia, Canada
BcRLR*: Burkholderia cepacia* Research Laboratory and Repository (John LiPuma, University of Michigan)
BDEC : Biodefense and Disease Ecology Center, University of North Arizona, USA

**Table S2. Comparison of in sillico and in vitro results of the distribution of the *hmqABCDEFG* operon**

| Strains | Prevalence of *hmqABCDEFG* operon (%) | |
|---|---|---|
| | Bioinformatics analysis [total genome sequences] | PCR analysis [total screened strains] |
| ***B. cepacia*** **(genomovar I)** | 23 [337] | 90 [31] |
| ***B. multivorans*** **(genomovar II)** | 0 [56] | 0 [45] |
| ***B. cenocepacia*** **(genomovar III)** | 0 [243] | 0 [72] |
| ***B. stabilis*** **(genomovar IV)** | - | 0 [21] |
| ***B. vietnamiensis*** **(genomovar V)** | 0 [41] | 13 [37] |
| ***B. dolosa*** **(genomovar VI)** | 0 [2] | 25 [4] |
| ***B. ambifaria*** **(genomovar VII)** | 50 [6] | 68 [32] |
| ***B. anthina*** **(genomovar VIII)** | 0 [8] | 25 [4] |
| ***B. pyrrocinia*** **(genomovar IX)** | 75 [4] | 78 [9] |
| ***B. ubonensis*** **(genomovar X)** | 97 [292] | 50 [2] |
| ***B. latens*** **(BCC1)** | 0 [2] | - |
| ***B. diffusa*** **(BCC 2)** | 0 [12] | 10 [10] |
| ***B. arboris*** **(BCC 3)** | - | 0 [4] |
| ***B. seminalis*** **(BCC 7)** | 0 [3] | 100 [1] |
| ***B. metallica*** **(BCC 8)** | 0 [1] | 25 [4] |
| ***B. lata*** **(group K)** | 50 [4] | 50 [4] |
| ***B. contaminans*** **(group K, BCCAT)** | 43 [7] | 82 [17] |
| ***B. pseudomultivorans*** | 0 [9] | - |
| ***B. stagnalis*** **(BCC B)** | 98 [64] | 100 [3] |
| ***B. territorii*** **(BCC I)** | 6 [33] | 100 [2] |
| ***B. paludis*** | - | - |
| **Other Bcc group** | 18 [59] | 45 [11] |
| **Total** | 35 [1257] | 30 [313] |

Not available data are represented by "-".

Kendall's rank test with a p-value of 0.01958

**Table S3. Study of the presence of the third chromosome in Bcc strains carrying or not the *hmqABCDEFG* operon.**

| Strains | # of chromosomes | Presence of the *hmqABCDEFG* operon |
|---|---|---|
| Burkholderia ambifaria AMMD | 3 | + |
| Burkholderia ambifaria MC40-6 | 3 | + |
| Burkholderia cenocepacia 842 | 3 | - |
| Burkholderia cenocepacia 895 | 2 | - |
| Burkholderia cenocepacia AU 1054 | 3 | - |
| Burkholderia cenocepacia CR318 | 3 | - |
| Burkholderia cenocepacia DDS 22E-1 | 3 | - |
| Burkholderia cenocepacia DWS 37E-2 | 3 | - |
| Burkholderia cenocepacia H111 | 3 | - |
| Burkholderia cenocepacia HI2424 | 3 | - |
| Burkholderia cenocepacia J2315 | 3 | - |
| Burkholderia cenocepacia MC0-3 | 3 | - |
| Burkholderia cenocepacia MSMB384WGS | 3 | - |
| Burkholderia cenocepacia VC12308 | 3 | - |
| Burkholderia cenocepacia VC12802 | 2 | - |
| Burkholderia cenocepacia VC7848 | 1 | - |
| Burkholderia cepacia ATCC 25416 | 3 | + |
| Burkholderia cepacia DDS 7H-2 | 3 | - |
| Burkholderia cepacia FDAARGOS_345 | 3 | + |
| Burkholderia cepacia FDAARGOS_388 | 3 | + |
| Burkholderia cepacia GG4 | 2 | - |
| Burkholderia cepacia INT3-BP177 | 2 | - |
| Burkholderia cepacia JBK9 | 3 | - |
| Burkholderia cepacia LO6 | 1 | - |
| Burkholderia cepacia MSMB1184WGS | 3 | + (c2) |
| Burkholderia contaminans MS14 | 3 | + |
| Burkholderia diffusa RF2-non-BP9 | 3 | - |
| Burkholderia dolosa AU0158 | 3 | - |
| Burkholderia lata 383 | 3 | - |
| Burkholderia lata FL-7-5-30-S1-D0 | 3 | + |
| Burkholderia latens AU17928 | 3 | - |
| Burkholderia metallica FL-6-5-30-S1-D7 | 3 | - |
| Burkholderia multivorans ATCC 17616 | 3 | - |
| Burkholderia multivorans ATCC 17616 | 3 | - |
| Burkholderia multivorans ATCC BAA-247 | 3 | - |
| Burkholderia multivorans AU1185 | 3 | - |
| Burkholderia multivorans MSMB1640WGS | 3 | - |
| Burkholderia pyrrocinia DSM 10685 | 3 | + |
| Burkholderia seminalis FL-5-4-10-S1-D7 | 3 | - |
| Burkholderia stabilis ATCC BAA-67 | 3 | - |
| Burkholderia stabilis FERMP-21014 | 3 | - |
| Burkholderia stagnalis MSMB735WGS | 3 | - |
| Burkholderia territorii RF8-non-BP5 | 3 | - |
| Burkholderia ubonensis MSMB0783 | 3 | - |
| Burkholderia ubonensis MSMB1189WGS | 3 | + |
| Burkholderia ubonensis MSMB1471WGS | 2 | + |
| Burkholderia ubonensis MSMB2035 | 3 | + |
| Burkholderia ubonensis MSMB22 | 3 | + |
| Burkholderia ubonensis RF23-BP41 | 3 | + |
| Burkholderia vietnamiensis AU1233 | 2 | - |
| Burkholderia vietnamiensis FL-2-3-30-S1-D0 | 3 | - |
| Burkholderia vietnamiensis G4 | 3 | - |
| Burkholderia vietnamiensis HI2297 | 3 | - |
| Burkholderia vietnamiensis LMG 10929 | 3 | - |
| Burkholderia vietnamiensis MSMB608WGS | 3 | - |

**Table S4. Quantification of the production of HMAQs for the Bcc strains which do not carry the *hmqABCDEFG* operon in their genome**

| Strains | Type | hmqA | hmqG | HMAQ_prod (TSB) |
|---|---|---|---|---|
| *B. ambifaria* CEP0516 | Clinical | - | - | - |
| *B. ambifaria* HI3590 | Environmental | - | - | - |
| *B. ambifaria* HI3687 | Environmental | - | - | - |
| *B. ambifaria* IOP40-10 | Environmental | - | - | - |
| *B. ambifaria* LMG17828 | Environmental | - | - | - |
| *B. ambifaria* PHP7 | Environmental | - | - | - |
| *B. anthina* VC16083 | Clinical | - | - | - |
| *B. arboris* VC8833 | Clinical | - | - | - |
| *B. cenocepacia* CEP024 | Clinical | - | - | - |
| *B. cenocepacia* CEP0511 | Clinical | - | - | - |
| *B. cenocepacia* CEP0565 | Clinical | - | - | - |
| *B. cenocepacia* IIIA VC16156 | Clinical | - | - | - |
| *B. cenocepacia* IIIB VC11311 | Clinical | - | - | - |
| *B. cenocepacia* IIIB VC15122 | Clinical | - | - | - |
| *B. cenocepacia* K56-2 | Clinical | - | - | - |
| *B. cenocepacia* LMG19240 | Environmental | - | - | - |
| *B. multivorans* CEP0781 | Clinical | - | - | - |
| *B. multivorans* LMG17588 | Environmental | - | - | - |
| *B. multivorans* VC14090 | Clinical | - | - | - |
| *B. multivorans* VC16759 | Clinical | - | - | - |
| *B. pyrrocinia* LMG21824 | Clinical | - | - | - |
| *B. stabilis* LMG18870 | Clinical | - | - | - |
| *B. stabilis* VC6749 | Clinical | - | - | - |
| *B. stabilis* VC6753 | Clinical | - | - | - |
| *B. ubonensis* LMG20358 | Environmental | - | - | - |
| B. vietnamiensis G4 | Environmental | - | - | - |
| *B. vietnamiensis* VC13984 | Clinical | - | - | - |
| *B. vietnamiensis* VC18210 | Clinical | - | - | - |
| *B. vietnamiensis* VC18712 | Clinical | - | - | - |
| *B. vietnamiensis* VC18844 | Clinical | - | - | - |
| *B. vietnamiensis* VC2824 | Clinical | - | - | - |

Table S5. HMAQs and HAQs produce by Bcc.

HMAQs production (mg/L/OD600nm)

| Strains | HMAQ-C7:2' Average | HMAQ-C7:2' Stdev | HMAQ-C9:2' Average | HMAQ-C9:2' Stdev | HMAQ-C5 Average | HMAQ-C5 Stdev | HHQ-C7 Average | HHQ-C7 Stdev | HMAQ-C6 Average | HMAQ-C6 Stdev | HMAQ-C7 Average | HMAQ-C7 Stdev |
|---|---|---|---|---|---|---|---|---|---|---|---|---|
| ***B. ambifaria* AU0212** | 8.7E-02 | 6.7E-02 | 4.1E-02 | 3.1E-02 | 0.0E+00 | 0.0E+00 | 0.0E+00 | 0.0E+00 | 0.0E+00 | 0.0E+00 | 0.0E+00 | 0.0E+00 |
| ***B. ambifaria* AU4157** | 1.7E+00 | 1.4E+00 | 1.3E+00 | 1.0E+00 | 5.8E-04 | 5.1E-04 | 0.0E+00 | 0.0E+00 | 0.0E+00 | 0.0E+00 | 4.5E-03 | 4.0E-03 |
| ***B. ambifaria* CEP0617** | 4.7E+00 | 4.9E-01 | 4.4E+00 | 3.5E-01 | 1.8E-03 | 2.5E-04 | 5.6E-03 | 4.4E-04 | 1.7E-03 | 2.3E-04 | 4.1E-02 | 6.1E-03 |
| ***B. ambifaria* CEP0958** | 2.9E+00 | 5.8E-02 | 2.8E+00 | 1.5E-01 | 1.3E-03 | 0.0E+00 | 1.5E-03 | 2.3E-04 | 0.0E+00 | 0.0E+00 | 8.5E-03 | 2.1E-04 |
| ***B. ambafaria* CEP0990** | 2.3E+00 | 1.1E+00 | 1.8E+00 | 9.2E-01 | 2.7E-03 | 1.4E-03 | 2.0E-03 | 1.5E-04 | 1.6E-03 | 5.8E-04 | 1.2E-02 | 6.0E-03 |
| ***B. ambifaria* CEP0996** | 2.6E+00 | 1.3E+00 | 1.4E+00 | 5.8E-01 | 1.3E-03 | 1.4E-03 | 2.8E-03 | 2.1E-04 | 0.0E+00 | 0.0E+00 | 6.0E-03 | 3.5E-03 |
| ***B. ambifaria* ES0020** | 2.1E+00 | 3.5E-01 | 1.8E+00 | 3.2E-01 | 9.9E-04 | 4.7E-04 | 1.0E-02 | 2.8E-03 | 3.7E-05 | 2.7E-05 | 3.6E-02 | 7.2E-03 |
| ***B. ambifaria* HI2468** | 6.0E-01 | 2.5E-02 | 4.2E-01 | 4.6E-02 | 2.6E-04 | 3.5E-05 | 1.7E-03 | 1.5E-04 | 3.3E-04 | 2.0E-04 | 6.2E-03 | 7.1E-04 |
| ***B. ambifaria* HI2482** | 7.0E-01 | 7.8E-02 | 4.1E-01 | 3.2E-02 | 8.1E-04 | 1.4E-04 | 3.3E-03 | 4.0E-04 | 5.6E-04 | 3.8E-04 | 6.3E-03 | 7.0E-04 |
| ***B. ambifaria* HI2626** | 8.6E-01 | 7.1E-02 | 6.1E-01 | 4.4E-02 | 9.0E-04 | 8.5E-05 | 3.0E-03 | 6.1E-04 | 2.7E-04 | 2.7E-04 | 7.3E-03 | 6.1E-04 |
| ***B. ambifaria* HI2672** | 1.1E+00 | 5.8E-02 | 8.1E-01 | 1.5E-02 | 1.3E-03 | 2.1E-04 | 2.1E-03 | 1.2E-04 | 4.0E-04 | 3.5E-04 | 1.2E-02 | 5.8E-04 |
| ***B. ambifaria* HI3709** | 4.3E-01 | 3.3E-01 | 3.1E-01 | 2.8E-01 | 5.2E-04 | 4.3E-04 | 1.9E-03 | 1.4E-03 | 1.8E-03 | 9.9E-04 | 6.9E-03 | 4.6E-03 |
| ***B. ambifaria* HI3738** | 9.3E-01 | 6.6E-01 | 6.1E-01 | 4.8E-01 | 1.3E-03 | 9.8E-04 | 2.7E-03 | 1.3E-03 | 9.6E-04 | 3.0E-04 | 1.2E-02 | 8.3E-03 |
| ***B. ambifaria* HSJ1** | 2.5E+00 | 1.1E+00 | 2.4E+00 | 1.1E+00 | 4.0E-04 | 6.9E-04 | 1.4E-03 | 0.0E+00 | 0.0E+00 | 0.0E+00 | 1.0E-02 | 5.3E-03 |
| ***B. ambifaria* PC736** | 9.8E-01 | 2.3E-01 | 6.1E-01 | 1.8E-01 | 1.0E-03 | 3.2E-04 | 2.1E-03 | 4.2E-04 | 4.5E-04 | 3.3E-04 | 6.0E-03 | 1.9E-03 |
| ***B. ambifaria* VC15422** | 5.5E-01 | 6.5E-01 | 3.8E-01 | 4.9E-01 | 0.0E+00 | 0.0E+00 | 2.1E-03 | 3.6E-03 | 3.3E-04 | 5.8E-04 | 1.5E-03 | 2.0E-03 |
| ***B. ambifaria* VC16196** | 1.1E+00 | 1.8E-01 | 9.5E-01 | 5.2E-02 | 0.0E+00 | 0.0E+00 | 2.1E-03 | 5.2E-04 | 2.0E-04 | 3.5E-04 | 2.0E-03 | 1.2E-04 |
| ***B. cepacia* BTS13** | 2.8E-01 | 4.3E-01 | 1.3E-01 | 1.8E-01 | 0.0E+00 | 0.0E+00 | 3.2E-04 | 5.5E-04 | 0.0E+00 | 0.0E+00 | 2.2E-03 | 3.9E-03 |
| ***B. cepacia* HI2430** | 3.2E+00 | 3.1E-01 | 2.3E+00 | 3.0E-01 | 7.8E-03 | 4.0E-04 | 1.9E-03 | 3.2E-04 | 2.8E-03 | 2.4E-03 | 2.1E-02 | 2.6E-03 |
| ***B. cepacia* HI2563** | 1.2E+00 | 3.5E-01 | 7.3E-01 | 2.3E-01 | 5.4E-03 | 1.0E-03 | 3.7E-03 | 1.8E-03 | 9.7E-05 | 1.4E-04 | 2.6E-02 | 8.4E-03 |
| ***B. cepacia* HI2578** | 5.5E-01 | 3.0E-01 | 2.6E-01 | 2.3E-01 | 1.0E-03 | 8.5E-04 | 7.1E-04 | 3.2E-04 | 0.0E+00 | 0.0E+00 | 7.9E-03 | 3.8E-03 |
| ***B. cepacia* HI2615** | 1.0E+00 | 5.8E-01 | 6.9E-01 | 4.4E-01 | 4.1E-03 | 2.1E-03 | 7.8E-03 | 3.1E-03 | 1.0E-03 | 6.9E-04 | 3.1E-02 | 1.5E-02 |
| ***B. cepacia* HI2671** | 4.9E-01 | 2.5E-01 | 2.8E-01 | 1.7E-01 | 7.9E-04 | 6.2E-04 | 3.1E-04 | 2.2E-04 | 5.2E-04 | 4.0E-04 | 2.2E-03 | 1.3E-03 |
| ***B. cepacia* HI2741** | 3.6E+00 | 7.6E-01 | 2.1E+00 | 5.0E-01 | 8.1E-03 | 2.9E-03 | 1.8E-03 | 7.0E-04 | 1.2E-04 | 1.8E-04 | 2.9E-02 | 9.8E-03 |
| ***B. cepacia* HI3312** | 6.3E-01 | 7.1E-02 | 4.1E-01 | 5.6E-02 | 1.4E-03 | 1.5E-04 | 2.8E-04 | 5.6E-05 | 3.7E-04 | 1.1E-04 | 5.0E-03 | 5.9E-04 |
| ***B. cepacia* HI3551** | 7.6E-01 | 3.2E-01 | 5.1E-01 | 2.6E-01 | 1.5E-03 | 7.0E-04 | 6.9E-04 | 2.5E-04 | 1.7E-03 | 1.3E-03 | 6.7E-03 | 2.9E-03 |
| ***B. cepacia* HI3708** | 1.3E+00 | 1.7E+00 | 6.1E-01 | 8.6E-01 | 5.0E-04 | 8.7E-04 | 5.3E-04 | 9.2E-04 | 2.7E-02 | 3.0E-02 | 1.3E-02 | 1.8E-02 |
| ***B. cepacia* HI3895** | 4.2E-01 | 0.0E+00 | 2.2E-01 | 1.7E-02 | 9.4E-04 | 9.8E-05 | 7.1E-04 | 6.1E-05 | 6.9E-04 | 6.0E-04 | 5.4E-03 | 2.5E-04 |
| ***B. cepacia* VC13132** | 1.9E-01 | 1.2E-01 | 1.3E-01 | 7.2E-02 | 0.0E+00 | 0.0E+00 | 0.0E+00 | 0.0E+00 | 0.0E+00 | 0.0E+00 | 6.3E-05 | 1.1E-04 |
| ***B. cepacia* VC13394** | 1.1E+00 | 1.2E-01 | 9.1E-01 | 9.0E-02 | 1.3E-03 | 2.1E-04 | 2.7E-03 | 5.5E-04 | 7.1E-04 | 1.6E-04 | 2.0E-03 | 2.6E-04 |
| ***B. cepacia* VC14106** | 8.9E-01 | 1.1E-01 | 7.8E-01 | 1.1E-01 | 5.8E-03 | 5.1E-03 | 0.0E+00 | 0.0E+00 | 2.3E-03 | 4.0E-03 | 4.0E-02 | 1.9E-02 |
| ***B. cepacia* VC14457** | 5.3E-01 | 6.0E-02 | 3.6E-01 | 2.0E-02 | 0.0E+00 | 0.0E+00 | 0.0E+00 | 0.0E+00 | 0.0E+00 | 0.0E+00 | 0.0E+00 | 0.0E+00 |
| ***B. cepacia* VC17333** | 5.1E-01 | 1.2E-01 | 4.8E-01 | 1.7E-01 | 0.0E+00 | 0.0E+00 | 6.5E-04 | 7.1E-04 | 0.0E+00 | 0.0E+00 | 5.2E-04 | 4.8E-04 |
| ***B. cepacia* VC17746** | 3.7E-01 | 3.3E-01 | 2.6E-01 | 2.3E-01 | 0.0E+00 | 0.0E+00 | 1.7E-04 | 2.9E-04 | 0.0E+00 | 0.0E+00 | 2.0E-04 | 3.5E-04 |
| ***B. cepacia* VC17928** | 4.5E-01 | 3.8E-01 | 4.2E-01 | 4.1E-01 | 3.3E-04 | 5.8E-04 | 9.3E-04 | 1.6E-03 | 2.3E-04 | 4.0E-04 | 5.3E-04 | 9.2E-04 |
| ***B. cepacia* VC18315** | 2.8E+00 | 1.5E-01 | 1.6E+00 | 5.8E-02 | 2.0E-03 | 3.2E-03 | 4.6E-04 | 5.1E-04 | 7.7E-04 | 1.3E-03 | 8.1E-03 | 8.2E-03 |
| ***B. cepacia* VC18839** | 1.1E-01 | 2.9E-02 | 4.8E-02 | 4.4E-02 | 0.0E+00 | 0.0E+00 | 0.0E+00 | 0.0E+00 | 0.0E+00 | 0.0E+00 | 1.4E-03 | 1.2E-03 |
| ***B. cepacia* VC18842** | 1.1E+00 | 7.7E-01 | 8.6E-01 | 6.0E-01 | 1.2E-03 | 1.1E-03 | 2.4E-03 | 2.1E-03 | 5.1E-04 | 4.5E-04 | 2.1E-03 | 1.8E-03 |
| ***B. cepacia* VC19276** | 4.2E-01 | 2.3E-01 | 3.4E-01 | 2.9E-01 | 0.0E+00 | 0.0E+00 | 4.5E-04 | 5.0E-04 | 0.0E+00 | 0.0E+00 | 5.3E-04 | 9.2E-04 |
| ***B. cepacia* VC9490** | 9.1E-01 | 1.9E-01 | 7.0E-01 | 7.5E-02 | 1.1E-03 | 2.6E-04 | 2.0E-03 | 6.8E-04 | 2.6E-04 | 4.4E-04 | 1.9E-03 | 5.2E-04 |
| ***B. contaminans* FFH2055** | 7.5E-01 | 4.8E-01 | 4.6E-01 | 4.1E-01 | 2.0E-04 | 3.5E-04 | 1.1E-03 | 3.2E-04 | 0.0E+00 | 0.0E+00 | 6.0E-03 | 3.3E-03 |
| ***B. contaminans* HI3570** | 7.5E-01 | 2.0E-01 | 2.9E-01 | 2.6E-01 | 1.0E-03 | 8.8E-04 | 6.9E-04 | 1.9E-04 | 2.0E-04 | 1.6E-04 | 9.3E-03 | 2.5E-03 |
| ***B. contaminans* HI3852** | 1.3E-01 | 7.7E-02 | 0.0E+00 | 0.0E+00 | 3.2E-05 | 3.4E-05 | 1.0E-04 | 2.3E-05 | 6.7E-04 | 5.8E-04 | 1.6E-03 | 6.6E-04 |
| ***B. contaminans* HI4067** | 4.6E-01 | 4.6E-02 | 0.0E+00 | 0.0E+00 | 3.5E-04 | 7.5E-05 | 5.4E-04 | 5.3E-05 | 5.4E-04 | 9.2E-04 | 6.2E-03 | 9.5E-04 |
| ***B. contaminans* HI4232** | 4.6E-01 | 8.6E-02 | 6.3E-02 | 1.1E-01 | 4.3E-04 | 7.8E-05 | 6.6E-04 | 4.7E-04 | 4.7E-04 | 3.0E-04 | 6.7E-03 | 1.3E-03 |
| ***B. contaminans* VC14347** | 1.5E-01 | 5.2E-02 | 0.0E+00 | 0.0E+00 | 0.0E+00 | 0.0E+00 | 0.0E+00 | 0.0E+00 | 0.0E+00 | 0.0E+00 | 0.0E+00 | 0.0E+00 |
| ***B. contaminans* VC16087** | 1.6E-01 | 2.9E-02 | 4.0E-02 | 6.9E-02 | 0.0E+00 | 0.0E+00 | 0.0E+00 | 0.0E+00 | 0.0E+00 | 0.0E+00 | 3.8E-03 | 1.4E-03 |
| ***B. contaminans* VC16087** | 2.0E-01 | 1.8E-01 | 0.0E+00 | 0.0E+00 | 0.0E+00 | 0.0E+00 | 0.0E+00 | 0.0E+00 | 0.0E+00 | 0.0E+00 | 0.0E+00 | 0.0E+00 |
| ***B. contaminans* VC16848-b** | 5.0E-01 | 9.2E-02 | 1.1E-02 | 1.2E-02 | 1.2E-04 | 2.1E-04 | 4.3E-04 | 4.8E-04 | 0.0E+00 | 0.0E+00 | 5.1E-04 | 5.0E-04 |
| ***B. contaminans* VC19056** | 4.5E-01 | 4.5E-02 | 7.7E-03 | 1.3E-02 | 7.7E-05 | 1.3E-04 | 1.1E-04 | 2.0E-04 | 0.0E+00 | 0.0E+00 | 5.4E-04 | 9.3E-05 |
| ***B. contaminans* VC19124** | 3.5E-01 | 1.5E-01 | 0.0E+00 | 0.0E+00 | 0.0E+00 | 0.0E+00 | 0.0E+00 | 0.0E+00 | 0.0E+00 | 0.0E+00 | 1.6E-02 | 6.4E-03 |
| ***B. contaminans* VC9624** | 1.5E-01 | 1.7E-01 | 2.1E-02 | 2.1E-02 | 2.6E-02 | 2.3E-02 | 5.9E-03 | 4.9E-03 | 5.2E-03 | 4.6E-03 | 7.0E-02 | 6.0E-02 |
| ***B. lata* BC01** | 2.2E+00 | 3.6E-01 | 2.9E-02 | 5.9E-03 | 2.5E-03 | 4.6E-04 | 1.1E-02 | 5.8E-04 | 9.0E-04 | 1.9E-04 | 5.0E-02 | 8.7E-03 |
| ***B. pyrrocinia BC02*** | 1.8E+00 | 6.4E-01 | 2.8E-02 | 1.2E-02 | 1.7E-03 | 7.6E-04 | 1.4E-02 | 3.5E-03 | 2.2E-05 | 2.2E-05 | 4.8E-02 | 1.5E-02 |
| ***B. pyrrocinia* CH-67** | 2.0E-01 | 1.0E-02 | 1.3E+00 | 1.2E-01 | 0.0E+00 | 0.0E+00 | 2.2E-03 | 3.5E-04 | 0.0E+00 | 0.0E+00 | 4.5E-03 | 2.6E-04 |
| ***B. pyrrocinia* HI2575** | 9.0E-01 | 2.1E-01 | 8.2E-01 | 2.5E-01 | 1.8E-03 | 9.5E-04 | 3.5E-03 | 5.1E-04 | 9.4E-04 | 1.4E-04 | 1.6E-02 | 4.7E-03 |
| ***B. pyrrocinia* HI2690** | 7.2E-01 | 2.5E-01 | 6.8E-01 | 2.8E-01 | 1.0E-03 | 1.5E-04 | 3.8E-03 | 9.0E-04 | 2.0E-04 | 3.4E-04 | 7.4E-03 | 3.2E-03 |
| ***B. pyrrocinia* HI3892** | 7.0E-01 | 1.5E-01 | 6.0E-01 | 1.7E-01 | 9.6E-04 | 2.4E-04 | 2.8E-03 | 6.6E-04 | 5.2E-04 | 1.3E-04 | 1.1E-02 | 3.0E-03 |
| ***B. stagnalis* MSMB1956WGS** | 2.9E-03 | 3.5E-04 | 1.4E-02 | 2.0E-03 | 0.0E+00 | 0.0E+00 | 0.0E+00 | 0.0E+00 | 0.0E+00 | 0.0E+00 | 0.0E+00 | 0.0E+00 |
| ***B. territorii* MSMB1301WGS** | 1.2E+00 | 5.3E-01 | 7.6E-01 | 2.2E-01 | 4.7E-04 | 8.1E-04 | 1.1E-03 | 1.2E-04 | 2.5E-04 | 4.3E-04 | 1.0E-02 | 7.7E-03 |
| ***B. territorii* MSMB1502WGS** | 5.1E-01 | 1.7E-01 | 3.1E-01 | 1.7E-01 | 0.0E+00 | 0.0E+00 | 1.3E-04 | 2.2E-04 | 0.0E+00 | 0.0E+00 | 2.5E-03 | 1.1E-03 |
| ***B. vietnamiensis* HI3392** | 7.1E-02 | 2.4E-02 | 4.3E-02 | 7.9E-03 | 9.0E-05 | 5.8E-05 | 2.0E-04 | 1.8E-04 | 2.4E-04 | 2.5E-05 | 7.9E-04 | 4.0E-04 |
| ***B. vietnamiensis* VC8245** | 5.6E-01 | 1.9E-01 | 5.0E-01 | 2.3E-01 | 5.7E-04 | 6.0E-04 | 9.7E-04 | 1.2E-03 | 2.2E-04 | 3.8E-04 | 8.6E-04 | 9.0E-04 |
| **Other Bcc BC06** | 8.8E-01 | 3.2E-01 | 0.0E+00 | 0.0E+00 | 1.3E-03 | 8.2E-04 | 1.1E-03 | 6.5E-04 | 5.0E-04 | 2.9E-04 | 8.3E-03 | 7.5E-03 |
| **Other Bcc BC13** | 1.9E+00 | 1.5E-01 | 3.4E-02 | 9.3E-03 | 4.0E-03 | 7.4E-04 | 1.1E-03 | 1.2E-04 | 8.8E-04 | 9.0E-04 | 2.5E-02 | 4.6E-03 |
| **Other Bcc ES0139** | 1.6E+00 | 5.7E-01 | 3.6E-02 | 1.6E-02 | 1.2E-03 | 4.6E-04 | 2.1E-03 | 7.6E-04 | 4.8E-04 | 5.4E-04 | 9.5E-03 | 4.3E-03 |
| **Other Bcc BC04** | 1.0E+00 | 3.6E-01 | 0.0E+00 | 0.0E+00 | 1.8E-03 | 5.5E-04 | 1.4E-03 | 4.0E-04 | 2.1E-03 | 1.8E-03 | 1.1E-02 | 4.6E-03 |
| **Other Bcc BC03** | 4.6E-02 | 3.1E-03 | 1.8E-01 | 1.2E-02 | 0.0E+00 | 0.0E+00 | 4.8E-05 | 3.0E-05 | 7.6E-05 | 7.1E-05 | 3.8E-04 | 5.1E-05 |
| **Other Bcc HI2701** | 1.2E+00 | 3.5E-01 | 1.2E+00 | 3.7E-01 | 1.6E-03 | 4.5E-04 | 2.3E-03 | 7.4E-04 | 1.9E-04 | 2.2E-04 | 1.5E-02 | 5.2E-03 |

| Strains | HHQ-C9:2' Average | HHQ-C9:2' Stdev | HMAQ-C8:2' Average | HMAQ-C8:2' Stdev | HHQ-C9 Average | HHQ-C9 Stdev | HMAQ-C8 Average | HMAQ-C8 Stdev | HMAQ-C9 Average | HMAQ-C9 Stdev | HMAQ-C10:2' Average | HMAQ-C10:2' Stdev | HMAQ-C10 Average | HMAQ-C10 Stdev | HMAQ-C11 Average | HMAQ-C11 Stdev |
|---|---|---|---|---|---|---|---|---|---|---|---|---|---|---|---|---|
| ***B. ambifaria* AU0212** | 0.0E+00 | 0.0E+00 | 0.0E+00 | 0.0E+00 | 0.0E+00 | 0.0E+00 | 0.0E+00 | 0.0E+00 | 0.0E+00 | 0.0E+00 | 0.0E+00 | 0.0E+00 | 0.0E+00 | 0.0E+00 | 0.0E+00 | 0.0E+00 |
| ***B. ambifaria* AU4157** | 3.7E-03 | 3.2E-03 | 0.0E+00 | 0.0E+00 | 4.6E-03 | 4.0E-03 | 4.6E-03 | 4.0E-03 | 0.0E+00 | 0.0E+00 | 0.0E+00 | 0.0E+00 | 0.0E+00 | 0.0E+00 | 0.0E+00 | 0.0E+00 |
| ***B. ambifaria* CEP0617** | 2.9E-02 | 3.6E-03 | 2.4E-03 | 2.3E-04 | 4.0E-02 | 3.2E-03 | 4.0E-02 | 3.2E-03 | 0.0E+00 | 0.0E+00 | 0.0E+00 | 0.0E+00 | 0.0E+00 | 0.0E+00 | 0.0E+00 | 0.0E+00 |
| ***B. ambifaria* CEP0958** | 2.2E-02 | 1.7E-03 | 6.0E-04 | 5.2E-04 | 1.3E-02 | 5.8E-04 | 1.3E-02 | 5.8E-04 | 0.0E+00 | 0.0E+00 | 0.0E+00 | 0.0E+00 | 1.0E-03 | 8.1E-05 | 0.0E+00 | 0.0E+00 |
| ***B. ambafaria* CEP0990** | 1.0E-02 | 8.5E-04 | 0.0E+00 | 0.0E+00 | 9.4E-03 | 4.0E-03 | 9.4E-03 | 4.0E-03 | 0.0E+00 | 0.0E+00 | 0.0E+00 | 0.0E+00 | 0.0E+00 | 0.0E+00 | 0.0E+00 | 0.0E+00 |
| ***B. ambifaria* CEP0996** | 1.9E-02 | 2.6E-03 | 0.0E+00 | 0.0E+00 | 3.7E-03 | 1.7E-03 | 3.7E-03 | 1.7E-03 | 0.0E+00 | 0.0E+00 | 0.0E+00 | 0.0E+00 | 0.0E+00 | 0.0E+00 | 0.0E+00 | 0.0E+00 |
| ***B. ambifaria* ES0020** | 6.5E-02 | 4.6E-03 | 4.5E-03 | 3.5E-04 | 0.0E+00 | 0.0E+00 | 7.7E-02 | 1.8E-02 | 1.0E-03 | 1.5E-03 | 5.3E-04 | 2.0E-04 | 4.0E-03 | 1.5E-03 | 4.1E-03 | 1.6E-03 |
| ***B. ambifaria* HI2468** | 1.7E-02 | 1.7E-03 | 5.7E-04 | 7.2E-05 | 0.0E+00 | 0.0E+00 | 7.4E-03 | 1.0E-03 | 1.1E-05 | 1.2E-05 | 1.1E-04 | 1.8E-05 | 4.1E-04 | 2.0E-05 | 3.4E-04 | 3.2E-05 |
| ***B. ambifaria* HI2482** | 2.0E-02 | 1.2E-03 | 6.7E-04 | 7.0E-05 | 0.0E+00 | 0.0E+00 | 6.3E-03 | 6.8E-04 | 2.5E-05 | 1.4E-05 | 1.3E-04 | 0.0E+00 | 5.3E-04 | 8.9E-05 | 5.0E-04 | 4.6E-05 |
| ***B. ambifaria* HI2626** | 2.8E-02 | 4.9E-03 | 8.8E-04 | 1.1E-04 | 0.0E+00 | 0.0E+00 | 9.0E-03 | 7.0E-04 | 4.6E-05 | 2.5E-06 | 1.8E-04 | 2.0E-05 | 8.1E-04 | 6.9E-05 | 7.8E-04 | 9.7E-05 |
| ***B. ambifaria* HI2672** | 2.2E-02 | 2.5E-03 | 8.2E-04 | 6.1E-05 | 0.0E+00 | 0.0E+00 | 1.7E-02 | 5.8E-04 | 3.7E-05 | 3.4E-05 | 4.6E-04 | 4.5E-05 | 1.8E-03 | 1.5E-04 | 1.6E-03 | 5.8E-05 |
| ***B. ambifaria* HI3709** | 1.3E-02 | 9.1E-03 | 6.0E-04 | 4.7E-04 | 0.0E+00 | 0.0E+00 | 9.6E-03 | 9.2E-03 | 1.2E-05 | 1.3E-05 | 8.3E-05 | 1.3E-04 | 4.0E-04 | 6.0E-04 | 4.4E-04 | 6.6E-04 |
| ***B. ambifaria* HI3738** | 1.7E-02 | 1.1E-02 | 9.4E-04 | 6.8E-04 | 0.0E+00 | 0.0E+00 | 1.8E-02 | 1.5E-02 | 6.1E-05 | 4.6E-05 | 2.8E-04 | 2.7E-04 | 1.6E-03 | 1.6E-03 | 1.3E-03 | 1.2E-03 |
| ***B. ambifaria* HSJ1** | 1.5E-02 | 5.8E-04 | 0.0E+00 | 0.0E+00 | 1.1E-02 | 5.3E-03 | 1.1E-02 | 5.3E-03 | 0.0E+00 | 0.0E+00 | 0.0E+00 | 0.0E+00 | 0.0E+00 | 0.0E+00 | 0.0E+00 | 0.0E+00 |
| ***B. ambifaria* PC736** | 2.4E-02 | 5.6E-03 | 7.1E-04 | 1.5E-04 | 0.0E+00 | 0.0E+00 | 8.0E-03 | 2.8E-03 | 0.0E+00 | 0.0E+00 | 2.2E-04 | 2.6E-05 | 8.8E-04 | 1.8E-04 | 3.6E-04 | 9.2E-05 |
| ***B. ambifaria* VC15422** | 1.8E-02 | 3.0E-02 | 5.3E-04 | 9.2E-04 | 1.3E-04 | 2.3E-04 | 9.0E-04 | 1.6E-03 | 0.0E+00 | 0.0E+00 | 0.0E+00 | 0.0E+00 | 0.0E+00 | 0.0E+00 | 0.0E+00 | 0.0E+00 |
| ***B. ambifaria* VC16196** | 3.0E-02 | 2.3E-03 | 2.7E-04 | 4.7E-04 | 0.0E+00 | 0.0E+00 | 2.1E-03 | 3.5E-04 | 0.0E+00 | 0.0E+00 | 0.0E+00 | 0.0E+00 | 0.0E+00 | 0.0E+00 | 0.0E+00 | 0.0E+00 |
| ***B. cepacia* BTS13** | 2.5E-03 | 3.5E-03 | 0.0E+00 | 0.0E+00 | 0.0E+00 | 0.0E+00 | 1.2E-03 | 2.1E-03 | 0.0E+00 | 0.0E+00 | 0.0E+00 | 0.0E+00 | 0.0E+00 | 0.0E+00 | 0.0E+00 | 0.0E+00 |
| ***B. cepacia* HI2430** | 2.5E-02 | 4.0E-03 | 8.8E-04 | 1.9E-04 | 0.0E+00 | 0.0E+00 | 4.5E-02 | 7.5E-03 | 1.2E-04 | 1.6E-05 | 2.0E-03 | 4.6E-04 | 4.0E-03 | 9.7E-04 | 1.9E-03 | 4.6E-04 |
| ***B. cepacia* HI2563** | 1.3E-02 | 2.1E-03 | 1.0E-03 | 3.3E-04 | 0.0E+00 | 0.0E+00 | 3.4E-02 | 1.1E-02 | 3.0E-05 | 3.2E-05 | 3.1E-04 | 1.7E-04 | 1.5E-03 | 6.4E-04 | 8.8E-04 | 2.1E-04 |
| ***B. cepacia* HI2578** | 3.5E-03 | 2.5E-03 | 1.5E-04 | 4.5E-05 | 0.0E+00 | 0.0E+00 | 5.4E-03 | 4.4E-03 | 5.7E-06 | 9.8E-06 | 1.0E-04 | 1.2E-04 | 1.8E-04 | 1.7E-04 | 1.0E-04 | 8.3E-05 |
| ***B. cepacia* HI2615** | 2.4E-02 | 1.1E-02 | 2.2E-03 | 1.0E-03 | 0.0E+00 | 0.0E+00 | 4.1E-02 | 2.6E-02 | 0.0E+00 | 0.0E+00 | 2.3E-04 | 1.9E-04 | 1.6E-03 | 1.1E-03 | 9.3E-04 | 7.5E-04 |
| ***B. cepacia* HI2671** | 4.0E-03 | 2.9E-03 | 8.2E-05 | 6.9E-05 | 1.6E-05 | 2.8E-05 | 2.2E-03 | 1.8E-03 | 1.0E-05 | 8.5E-06 | 7.7E-05 | 5.5E-05 | 1.2E-04 | 1.3E-04 | 6.7E-05 | 8.1E-05 |
| ***B. cepacia* HI2741** | 1.6E-02 | 1.7E-03 | 7.1E-04 | 1.9E-04 | 0.0E+00 | 0.0E+00 | 4.0E-02 | 1.7E-02 | 1.5E-04 | 6.3E-05 | 1.2E-03 | 3.4E-04 | 2.3E-03 | 9.5E-04 | 9.4E-04 | 3.7E-04 |
| ***B. cepacia* HI3312** | 3.8E-03 | 6.1E-04 | 1.3E-04 | 1.0E-05 | 0.0E+00 | 0.0E+00 | 9.1E-03 | 1.8E-03 | 2.0E-05 | 1.8E-05 | 2.6E-04 | 7.2E-05 | 5.8E-04 | 1.3E-04 | 2.5E-04 | 8.4E-05 |
| ***B. cepacia* HI3551** | 5.2E-03 | 2.1E-03 | 1.3E-04 | 1.0E-04 | 0.0E+00 | 0.0E+00 | 5.2E-03 | 3.0E-03 | 2.6E-06 | 4.5E-06 | 2.3E-04 | 1.5E-04 | 2.5E-04 | 1.8E-04 | 1.6E-04 | 1.0E-04 |
| ***B. cepacia* HI3708** | 8.3E-03 | 8.4E-03 | 0.0E+00 | 0.0E+00 | 0.0E+00 | 0.0E+00 | 9.5E-03 | 1.2E-02 | 0.0E+00 | 0.0E+00 | 0.0E+00 | 0.0E+00 | 0.0E+00 | 0.0E+00 | 0.0E+00 | 0.0E+00 |
| ***B. cepacia* HI3895** | 3.2E-03 | 3.2E-04 | 1.2E-04 | 2.2E-05 | 0.0E+00 | 0.0E+00 | 3.2E-03 | 1.5E-04 | 5.7E-06 | 9.8E-06 | 7.5E-05 | 2.2E-05 | 1.2E-04 | 2.3E-05 | 4.9E-05 | 1.7E-05 |
| ***B. cepacia* VC13132** | 1.8E-03 | 8.6E-04 | 0.0E+00 | 0.0E+00 | 5.7E-05 | 9.8E-05 | 0.0E+00 | 0.0E+00 | 0.0E+00 | 0.0E+00 | 0.0E+00 | 0.0E+00 | 0.0E+00 | 0.0E+00 | 0.0E+00 | 0.0E+00 |
| ***B. cepacia* VC13394** | 3.1E-02 | 3.0E-03 | 6.9E-04 | 7.8E-05 | 0.0E+00 | 0.0E+00 | 2.1E-03 | 2.1E-04 | 0.0E+00 | 0.0E+00 | 0.0E+00 | 0.0E+00 | 0.0E+00 | 0.0E+00 | 0.0E+00 | 0.0E+00 |
| ***B. cepacia* VC14106** | 1.7E-02 | 1.7E-03 | 5.2E-02 | 1.1E-02 | 0.0E+00 | 0.0E+00 | 0.0E+00 | 0.0E+00 | 7.3E-02 | 1.3E-02 | 0.0E+00 | 0.0E+00 | 7.3E-03 | 6.7E-04 | 0.0E+00 | 0.0E+00 |
| ***B. cepacia* VC14457** | 7.3E-03 | 6.8E-04 | 0.0E+00 | 0.0E+00 | 0.0E+00 | 0.0E+00 | 0.0E+00 | 0.0E+00 | 0.0E+00 | 0.0E+00 | 0.0E+00 | 0.0E+00 | 0.0E+00 | 0.0E+00 | 0.0E+00 | 0.0E+00 |
| ***B. cepacia* VC17333** | 8.6E-03 | 3.2E-03 | 1.6E-04 | 2.7E-04 | 0.0E+00 | 0.0E+00 | 6.6E-04 | 5.8E-04 | 0.0E+00 | 0.0E+00 | 0.0E+00 | 0.0E+00 | 0.0E+00 | 0.0E+00 | 0.0E+00 | 0.0E+00 |
| ***B. cepacia* VC17746** | 5.5E-03 | 4.9E-03 | 0.0E+00 | 0.0E+00 | 0.0E+00 | 0.0E+00 | 2.1E-04 | 3.6E-04 | 0.0E+00 | 0.0E+00 | 0.0E+00 | 0.0E+00 | 0.0E+00 | 0.0E+00 | 0.0E+00 | 0.0E+00 |
| ***B. cepacia* VC17928** | 1.3E-02 | 1.5E-02 | 2.6E-04 | 4.5E-04 | 0.0E+00 | 0.0E+00 | 6.7E-04 | 1.2E-03 | 0.0E+00 | 0.0E+00 | 0.0E+00 | 0.0E+00 | 0.0E+00 | 0.0E+00 | 0.0E+00 | 0.0E+00 |
| ***B. cepacia* VC18315** | 1.3E-02 | 2.0E-02 | 1.7E-04 | 2.9E-04 | 9.7E-03 | 1.6E-02 | 9.9E-03 | 1.6E-02 | 2.4E-03 | 4.1E-03 | 2.9E-03 | 5.0E-03 | 2.4E-03 | 4.1E-03 | 1.1E-03 | 1.8E-03 |
| ***B. cepacia* VC18839** | 0.0E+00 | 0.0E+00 | 9.7E-04 | 1.7E-03 | 0.0E+00 | 0.0E+00 | 0.0E+00 | 0.0E+00 | 0.0E+00 | 0.0E+00 | 0.0E+00 | 0.0E+00 | 0.0E+00 | 0.0E+00 | 0.0E+00 | 0.0E+00 |
| ***B. cepacia* VC18842** | 2.8E-02 | 2.2E-02 | 3.0E-04 | 5.1E-04 | 0.0E+00 | 0.0E+00 | 1.6E-03 | 1.4E-03 | 0.0E+00 | 0.0E+00 | 0.0E+00 | 0.0E+00 | 0.0E+00 | 0.0E+00 | 0.0E+00 | 0.0E+00 |
| ***B. cepacia* VC19276** | 7.1E-03 | 6.9E-03 | 0.0E+00 | 0.0E+00 | 0.0E+00 | 0.0E+00 | 5.0E-04 | 8.7E-04 | 0.0E+00 | 0.0E+00 | 0.0E+00 | 0.0E+00 | 0.0E+00 | 0.0E+00 | 0.0E+00 | 0.0E+00 |
| ***B. cepacia* VC9490** | 1.9E-02 | 3.8E-03 | 2.2E-04 | 3.8E-04 | 0.0E+00 | 0.0E+00 | 1.1E-03 | 6.4E-04 | 0.0E+00 | 0.0E+00 | 0.0E+00 | 0.0E+00 | 0.0E+00 | 0.0E+00 | 0.0E+00 | 0.0E+00 |
| ***B. contaminans* FFH2055** | 4.3E-03 | 2.3E-04 | 1.7E-04 | 3.0E-04 | 0.0E+00 | 0.0E+00 | 3.7E-03 | 1.9E-03 | 0.0E+00 | 0.0E+00 | 0.0E+00 | 0.0E+00 | 0.0E+00 | 0.0E+00 | 0.0E+00 | 0.0E+00 |
| ***B. contaminans* HI3570** | 3.4E-03 | 2.9E-03 | 2.1E-04 | 9.0E-05 | 1.7E-06 | 3.0E-06 | 5.4E-03 | 4.9E-03 | 4.7E-06 | 8.1E-06 | 1.4E-04 | 1.3E-04 | 3.4E-04 | 3.5E-04 | 1.7E-04 | 1.5E-04 |
| ***B. contaminans* HI3852** | 2.3E-05 | 1.7E-05 | 1.6E-05 | 1.7E-05 | 1.7E-05 | 1.5E-05 | 3.2E-06 | 5.5E-06 | 0.0E+00 | 0.0E+00 | 0.0E+00 | 0.0E+00 | 0.0E+00 | 0.0E+00 | 0.0E+00 | 0.0E+00 |
| ***B. contaminans* HI4067** | 8.6E-05 | 1.2E-05 | 1.0E-04 | 5.7E-05 | 1.1E-05 | 4.2E-06 | 4.8E-05 | 1.7E-05 | 0.0E+00 | 0.0E+00 | 0.0E+00 | 0.0E+00 | 0.0E+00 | 0.0E+00 | 4.6E-06 | 4.7E-06 |
| ***B. contaminans* HI4232** | 1.4E-03 | 2.2E-03 | 1.1E-04 | 9.5E-05 | 1.3E-05 | 1.2E-05 | 1.4E-03 | 2.3E-03 | 0.0E+00 | 0.0E+00 | 1.2E-05 | 2.0E-05 | 8.0E-05 | 1.4E-04 | 6.0E-05 | 1.0E-04 |
| ***B. contaminans* VC14347** | 0.0E+00 | 0.0E+00 | 0.0E+00 | 0.0E+00 | 0.0E+00 | 0.0E+00 | 0.0E+00 | 0.0E+00 | 0.0E+00 | 0.0E+00 | 0.0E+00 | 0.0E+00 | 0.0E+00 | 0.0E+00 | 0.0E+00 | 0.0E+00 |
| ***B. contaminans* VC16087** | 0.0E+00 | 0.0E+00 | 3.1E-03 | 2.3E-04 | 0.0E+00 | 0.0E+00 | 0.0E+00 | 0.0E+00 | 1.2E-03 | 2.0E-03 | 0.0E+00 | 0.0E+00 | 0.0E+00 | 0.0E+00 | 0.0E+00 | 0.0E+00 |
| ***B. contaminans* VC16087** | 0.0E+00 | 0.0E+00 | 0.0E+00 | 0.0E+00 | 0.0E+00 | 0.0E+00 | 0.0E+00 | 0.0E+00 | 0.0E+00 | 0.0E+00 | 0.0E+00 | 0.0E+00 | 0.0E+00 | 0.0E+00 | 0.0E+00 | 0.0E+00 |
| ***B. contaminans* VC16848-b** | 5.7E-05 | 9.8E-05 | 0.0E+00 | 0.0E+00 | 9.3E-05 | 1.6E-04 | 0.0E+00 | 0.0E+00 | 0.0E+00 | 0.0E+00 | 0.0E+00 | 0.0E+00 | 0.0E+00 | 0.0E+00 | 0.0E+00 | 0.0E+00 |
| ***B. contaminans* VC19056** | 0.0E+00 | 0.0E+00 | 0.0E+00 | 0.0E+00 | 8.3E-05 | 1.4E-04 | 0.0E+00 | 0.0E+00 | 0.0E+00 | 0.0E+00 | 0.0E+00 | 0.0E+00 | 0.0E+00 | 0.0E+00 | 0.0E+00 | 0.0E+00 |
| ***B. contaminans* VC19124** | 0.0E+00 | 0.0E+00 | 0.0E+00 | 0.0E+00 | 0.0E+00 | 0.0E+00 | 0.0E+00 | 0.0E+00 | 0.0E+00 | 0.0E+00 | 0.0E+00 | 0.0E+00 | 0.0E+00 | 0.0E+00 | 0.0E+00 | 0.0E+00 |
| ***B. contaminans* VC9624** | 5.4E-02 | 4.7E-02 | 2.7E-03 | 2.4E-03 | 1.3E-01 | 1.1E-01 | 1.3E-01 | 1.1E-01 | 5.5E-03 | 9.1E-03 | 3.9E-03 | 3.6E-03 | 1.1E-02 | 9.2E-03 | 4.6E-03 | 4.1E-03 |
| ***B. lata* BC01** | 1.3E-03 | 1.7E-04 | 9.2E-03 | 1.6E-03 | 0.0E+00 | 0.0E+00 | 3.4E-03 | 7.5E-04 | 0.0E+00 | 0.0E+00 | 0.0E+00 | 0.0E+00 | 3.9E-05 | 1.2E-05 | 6.1E-05 | 5.6E-06 |
| ***B. pyrrocinia BC02*** | 6.4E-04 | 3.3E-04 | 3.9E-03 | 2.6E-03 | 0.0E+00 | 0.0E+00 | 1.6E-03 | 6.2E-04 | 0.0E+00 | 0.0E+00 | 0.0E+00 | 0.0E+00 | 1.9E-05 | 1.1E-05 | 4.5E-05 | 1.9E-05 |
| ***B. pyrrocinia* CH-67** | 1.0E-01 | 1.1E-02 | 4.5E-03 | 6.0E-04 | 0.0E+00 | 0.0E+00 | 2.9E-02 | 1.7E-03 | 0.0E+00 | 0.0E+00 | 0.0E+00 | 0.0E+00 | 0.0E+00 | 0.0E+00 | 0.0E+00 | 0.0E+00 |
| ***B. pyrrocinia* HI2575** | 3.1E-02 | 1.4E-02 | 2.0E-03 | 9.5E-04 | 0.0E+00 | 0.0E+00 | 4.3E-02 | 1.9E-02 | 3.2E-05 | 3.6E-05 | 1.5E-04 | 5.9E-05 | 1.6E-03 | 1.2E-03 | 3.3E-03 | 2.9E-03 |
| ***B. pyrrocinia* HI2690** | 4.7E-02 | 1.9E-02 | 3.1E-03 | 1.3E-03 | 0.0E+00 | 0.0E+00 | 3.5E-02 | 1.8E-02 | 1.7E-05 | 2.9E-05 | 1.3E-04 | 1.7E-05 | 2.0E-03 | 7.4E-04 | 1.5E-03 | 1.1E-03 |
| ***B. pyrrocinia* HI3892** | 1.9E-02 | 2.1E-03 | 1.2E-03 | 3.8E-04 | 0.0E+00 | 0.0E+00 | 1.9E-02 | 7.9E-03 | 2.6E-05 | 1.2E-05 | 4.4E-05 | 1.2E-05 | 5.4E-04 | 2.7E-04 | 2.3E-04 | 1.2E-04 |
| ***B. stagnalis* MSMB1956WGS** | 9.7E-05 | 1.7E-04 | 0.0E+00 | 0.0E+00 | 0.0E+00 | 0.0E+00 | 0.0E+00 | 0.0E+00 | 0.0E+00 | 0.0E+00 | 0.0E+00 | 0.0E+00 | 0.0E+00 | 0.0E+00 | 0.0E+00 | 0.0E+00 |
| ***B. territorii* MSMB1301WGS** | 5.6E-03 | 1.4E-03 | 0.0E+00 | 0.0E+00 | 0.0E+00 | 0.0E+00 | 7.2E-03 | 5.9E-03 | 0.0E+00 | 0.0E+00 | 0.0E+00 | 0.0E+00 | 0.0E+00 | 0.0E+00 | 0.0E+00 | 0.0E+00 |
| ***B. territorii* MSMB1502WGS** | 2.2E-03 | 2.5E-04 | 0.0E+00 | 0.0E+00 | 0.0E+00 | 0.0E+00 | 1.5E-03 | 7.6E-04 | 0.0E+00 | 0.0E+00 | 0.0E+00 | 0.0E+00 | 0.0E+00 | 0.0E+00 | 0.0E+00 | 0.0E+00 |
| ***B. vietnamiensis* HI3392** | 9.9E-04 | 4.5E-04 | 6.7E-05 | 5.6E-05 | 1.3E-05 | 1.2E-05 | 7.1E-04 | 3.2E-04 | 0.0E+00 | 0.0E+00 | 3.7E-06 | 2.9E-06 | 1.7E-05 | 1.1E-05 | 4.5E-06 | 5.1E-06 |
| ***B. vietnamiensis* VC8245** | 1.0E-02 | 7.0E-03 | 1.3E-04 | 2.3E-04 | 0.0E+00 | 0.0E+00 | 6.8E-04 | 7.6E-04 | 0.0E+00 | 0.0E+00 | 0.0E+00 | 0.0E+00 | 0.0E+00 | 0.0E+00 | 0.0E+00 | 0.0E+00 |
| **Other Bcc BC06** | 4.6E-04 | 1.6E-04 | 6.5E-04 | 9.9E-04 | 1.7E-05 | 1.5E-05 | 3.4E-04 | 3.7E-04 | 0.0E+00 | 0.0E+00 | 0.0E+00 | 0.0E+00 | 0.0E+00 | 0.0E+00 | 4.7E-06 | 8.1E-06 |
| **Other Bcc BC13** | 6.8E-04 | 1.4E-04 | 7.3E-04 | 1.8E-04 | 2.0E-05 | 3.4E-05 | 1.4E-03 | 3.6E-04 | 0.0E+00 | 0.0E+00 | 1.2E-05 | 1.6E-05 | 2.0E-06 | 3.0E-06 | 1.3E-05 | 6.0E-06 |
| **Other Bcc ES0139** | 8.9E-04 | 2.7E-04 | 2.2E-04 | 9.7E-05 | 1.8E-05 | 1.4E-05 | 3.6E-04 | 1.7E-04 | 0.0E+00 | 0.0E+00 | 0.0E+00 | 0.0E+00 | 0.0E+00 | 0.0E+00 | 0.0E+00 | 0.0E+00 |
| **Other Bcc BC04** | 3.8E-04 | 1.9E-04 | 3.1E-04 | 1.7E-04 | 2.4E-05 | 8.7E-06 | 2.3E-04 | 1.4E-04 | 0.0E+00 | 0.0E+00 | 0.0E+00 | 0.0E+00 | 3.3E-06 | 5.7E-06 | 1.5E-05 | 1.2E-05 |
| **Other Bcc BC03** | 3.7E-03 | 5.3E-04 | 9.9E-05 | 1.2E-06 | 0.0E+00 | 0.0E+00 | 2.6E-03 | 5.8E-05 | 3.7E-06 | 6.4E-06 | 6.8E-05 | 9.8E-06 | 6.3E-05 | 2.0E-05 | 6.2E-06 | 5.4E-06 |
| **Other Bcc HI2701** | 3.3E-02 | 9.0E-03 | 1.9E-03 | 6.2E-04 | 0.0E+00 | 0.0E+00 | 5.6E-02 | 1.9E-02 | 4.0E-05 | 6.9E-05 | 4.2E-04 | 1.2E-04 | 4.2E-03 | 1.4E-03 | 5.1E-03 | 1.9E-03 |

an heatmap was generated for each produced molecule

**Table S6. Average of HMAQ-C7:1 and HMAQ-C9:1 production in Bcc.**

| Strains | | HMAQ-C7:2' and HMAQ-C9:2' production by strain HMAQ-C7:2' Average | Stdev | HMAQ-C9:2' Average | Stdev | by species HMAQ-C7:2' Average | Stdev | HMAQ-C9:2' Average | Stdev | by type HMAQ-C7:2' Average | Stdev | HMAQ-C9:2' Average | Stdev |
|---|---|---|---|---|---|---|---|---|---|---|---|---|---|
| | *B. ambifaria* AU0212 | 0.09 | 0.07 | 0.04 | 0.03 | | | | | | | | |
| | *B. ambifaria* AU4157 | 1.73 | 1.37 | 1.26 | 1.04 | | | | | | | | |
| | *B. ambifaria* CEP0617 | 4.73 | 0.49 | 4.40 | 0.35 | | | | | | | | |
| | *B. ambifaria* CEP0958 | 2.87 | 0.06 | 2.83 | 0.15 | | | | | | | | |
| | *B. ambafaria* CEP0990 | 2.30 | 1.13 | 1.83 | 0.92 | 2.06 | 0.53 | 1.72 | 0.41 | | | | |
| | *B. ambifaria* CEP0996 | 2.63 | 1.31 | 1.41 | 0.58 | | | | | | | | |
| | *B. ambifaria* HSJ1 | 2.53 | 1.05 | 2.37 | 1.05 | | | | | | | | |
| | *B. ambifaria* VC15422 | 0.55 | 0.65 | 0.38 | 0.49 | | | | | | | | |
| | *B. ambifaria* VC16196 | 1.09 | 0.18 | 0.95 | 0.05 | | | | | | | | |
| | *B. cepacia* BTS13 | 0.28 | 0.43 | 0.13 | 0.18 | | | | | | | | |
| | *B. cepacia* VC13132 | 0.19 | 0.12 | 0.13 | 0.07 | | | | | | | | |
| | *B. cepacia* VC13394 | 1.06 | 0.12 | 0.91 | 0.09 | | | | | | | | |
| | *B. cepacia* VC14106 | 0.89 | 0.11 | 0.78 | 0.11 | | | | | | | | |
| | *B. cepacia* VC14457 | 0.53 | 0.06 | 0.36 | 0.02 | | | | | | | | |
| Clinical Bcc | *B. cepacia* VC17333 | 0.51 | 0.12 | 0.48 | 0.17 | | | | | 1.04 | 1.11 | 0.78 | 1.00 |
| | *B. cepacia* VC17746 | 0.37 | 0.33 | 0.26 | 0.23 | 0.74 | 0.71 | 0.54 | 0.42 | | | | |
| | *B. cepacia* VC17928 | 0.45 | 0.38 | 0.42 | 0.41 | | | | | | | | |
| | *B. cepacia* VC18315 | 2.83 | 0.15 | 1.57 | 0.06 | | | | | | | | |
| | *B. cepacia* VC18839 | 0.11 | 0.03 | 0.05 | 0.04 | | | | | | | | |
| | *B. cepacia* VC18842 | 1.05 | 0.77 | 0.86 | 0.60 | | | | | | | | |
| | *B. cepacia* VC19276 | 0.42 | 0.23 | 0.34 | 0.29 | | | | | | | | |
| | *B. cepacia* VC9490 | 0.91 | 0.19 | 0.70 | 0.08 | | | | | | | | |
| | *B. contaminans* FFH2055 | 0.75 | 0.48 | 0.46 | 0.41 | | | | | | | | |
| | *B. contaminans* VC14347 | 0.15 | 0.05 | 0.00 | 0.00 | | | | | | | | |
| | *B. contaminans* VC16087 | 0.16 | 0.03 | 0.04 | 0.07 | | | | | | | | |
| | *B. contaminans* VC16848-b | 0.50 | 0.09 | 0.01 | 0.01 | 0.36 | 0.23 | 0.08 | 0.17 | | | | |
| | *B. contaminans* VC19056 | 0.45 | 0.05 | 0.01 | 0.01 | | | | | | | | |
| | *B. contaminans* VC19124 | 0.35 | 0.15 | 0.00 | 0.00 | | | | | | | | |
| | *B. contaminans* VC9624 | 0.15 | 0.17 | 0.02 | 0.02 | | | | | | | | |
| | *B. vietnamiensis* VC8245 | 0.56 | 0.19 | 0.50 | 0.23 | 0.60 | 0.10 | 0.31 | 0.17 | | | | |
| | *B. ambifaria* ES0020 | 2.07 | 0.35 | 1.77 | 0.32 | | | | | | | | |
| | *B. ambifaria* HI2468 | 0.60 | 0.03 | 0.42 | 0.05 | | | | | | | | |
| | *B. ambifaria* HI2482 | 0.70 | 0.08 | 0.41 | 0.03 | | | | | | | | |
| | *B. ambifaria* HI2626 | 0.86 | 0.07 | 0.61 | 0.04 | 0.96 | 0.50 | 0.69 | 0.46 | | | | |
| | *B. ambifaria* HI2672 | 1.13 | 0.06 | 0.81 | 0.02 | | | | | | | | |
| | *B. ambifaria* HI3709 | 0.43 | 0.33 | 0.31 | 0.28 | | | | | | | | |
| | *B. ambifaria* HI3738 | 0.93 | 0.66 | 0.61 | 0.48 | | | | | | | | |
| | *B. ambifaria* PC736 | 0.98 | 0.23 | 0.61 | 0.18 | | | | | | | | |
| | *B. cepacia* HI2430 | 3.17 | 0.31 | 2.30 | 0.30 | | | | | | | | |
| | *B. cepacia* HI2563 | 1.17 | 0.35 | 0.73 | 0.23 | | | | | | | | |
| | *B. cepacia* HI2578 | 0.55 | 0.30 | 0.26 | 0.23 | | | | | | | | |
| | *B. cepacia* HI2615 | 1.04 | 0.58 | 0.69 | 0.44 | | | | | | | | |
| | *B. cepacia* HI2671 | 0.49 | 0.25 | 0.28 | 0.17 | 1.31 | 1.13 | 0.81 | 0.76 | | | | |
| | *B. cepacia* HI2741 | 3.57 | 0.76 | 2.13 | 0.50 | | | | | | | | |
| | *B. cepacia* HI3312 | 0.63 | 0.07 | 0.41 | 0.06 | | | | | | | | |
| | *B. cepacia* HI3551 | 0.76 | 0.32 | 0.51 | 0.26 | | | | | | | | |
| | *B. cepacia* HI3708 | 1.26 | 1.68 | 0.61 | 0.86 | | | | | | | | |
| | *B. cepacia* HI3895 | 0.42 | 0.00 | 0.22 | 0.02 | | | | | | | | |
| Environmental Bcc | *B. contaminans* HI3570 | 0.75 | 0.20 | 0.29 | 0.26 | | | | | 0.99 | 0.78 | 0.53 | 0.57 |
| | *B. contaminans* HI3852 | 0.13 | 0.08 | 0.00 | 0.00 | 0.45 | 0.25 | 0.09 | 0.14 | | | | |
| | *B. contaminans* HI4067 | 0.46 | 0.05 | 0.00 | 0.00 | | | | | | | | |
| | *B. contaminans* HI4232 | 0.46 | 0.09 | 0.06 | 0.11 | | | | | | | | |
| | *B. lata* BC01 | 2.20 | 0.36 | 0.03 | 0.01 | 2.20 | 0.36 | 0.03 | 0.01 | | | | |
| | *B. pyrrocinia* BC02 | 1.77 | 0.64 | 0.03 | 0.01 | | | | | | | | |
| | *B. pyrrocinia* CH-67 | 0.20 | 0.01 | 1.27 | 0.12 | | | | | | | | |
| | *B. pyrrocinia* HI2575 | 0.90 | 0.21 | 0.82 | 0.25 | 0.92 | 0.53 | 0.77 | 0.46 | | | | |
| | *B. pyrrocinia* HI2690 | 0.72 | 0.25 | 0.68 | 0.28 | | | | | | | | |
| | *B. pyrrocinia* HI2701 | 1.24 | 0.35 | 1.22 | 0.37 | | | | | | | | |
| | *B. pyrrocinia* HI3892 | 0.70 | 0.15 | 0.60 | 0.17 | | | | | | | | |
| | *B. stagnalis* MSMB1956WGS | 0.00 | 0.00 | 0.01 | 0.00 | 0.00 | 0.00 | 0.01 | 0.00 | | | | |
| | *B. territorii* MSMB1301WGS | 1.19 | 0.53 | 0.76 | 0.22 | 0.85 | 0.48 | 0.54 | 0.31 | | | | |
| | *B. territorii* MSMB1502WGS | 0.51 | 0.17 | 0.31 | 0.17 | | | | | | | | |
| | *B. vietnamiensis* HI3392 | 0.07 | 0.02 | 0.04 | 0.01 | 0.07 | 0.02 | 0.04 | 0.01 | | | | |
| | Other Bcc group 1 BC06 | 0.88 | 0.32 | 0.00 | 0.00 | | | | | | | | |
| | Other Bcc group 3 BC13 | 1.93 | 0.15 | 0.03 | 0.01 | | | | | | | | |
| | Other Bcc group 3 ES0139 | 1.57 | 0.57 | 0.04 | 0.02 | 1.09 | 0.72 | 0.05 | 0.07 | | | | |
| | Other Bcc group 4 BC04 | 1.03 | 0.36 | 0.00 | 0.00 | | | | | | | | |
| | Other Bcc group 5 BC03 | 0.05 | 0.00 | 0.18 | 0.01 | | | | | | | | |

**Table S7. Correlation data of the presence of the *hmqABCDEF* operon and the production of HMAQ in Bcc.**

| Species | *hmq operon* | HMAQs | Co-isolated with *Pseudomonas* +/- 7 days | *Pseudomonas* in the previous year | Source |
|---|---|---|---|---|---|
| ***B. cenocepacia*** **IIIA VC12308** | no | no | yes | yes | sputum |
| ***B. cenocepacia*** **IIIA VC15419** | no | no | no | no | sputum |
| ***B. cenocepacia*** **IIIA VC18585** | no | no | yes | no | respiratory |
| ***B. cenocepacia*** **IIIA VC18996** | no | no | no | no | sputum |
| ***B. cenocepacia*** **IIIA VC18999** | no | no | no | no | throat |
| ***B. cenocepacia*** **IIIA VC3917** | no | no | yes | yes | sputum |
| ***B. cenocepacia*** **IIIA VC6356** | no | no | no | yes | sputum |
| ***B. cenocepacia*** **IIIA VC6553** | no | no | no | yes | sputum |
| ***B. cenocepacia*** **IIIB VC11311** | no | no | yes | yes | sputum |
| ***B. cenocepacia*** **IIIB VC15122** | no | no | no | no | throat |
| ***B. cenocepacia*** **IIIB VC6598** | no | no | yes | yes | sputum |
| ***B. cenocepacia*** **IIIB VC7349** | no | no | no | yes | respiratory |
| ***B. cenocepacia*** **IIIB VC7849** | no | no | no | yes | sputum |
| ***B. cenocepacia*** **IIIB VC7911** | no | no | no | no | respiratory |
| ***B. cenocepacia*** **IIIB VC8340** | no | no | yes | yes | respiratory |
| ***B. cepacia*** **VC18315** | yes | yes | no | yes | respiratory |
| ***B. cepacia*** **VC9490** | yes | yes | yes | no | sputum |
| ***B. contaminans*** **VC14347** | yes | yes | yes | yes | respiratory |
| ***B. contaminans*** **VC15406** | yes | no | no | yes | sinus |
| ***B. contaminans*** **VC16087** | yes | yes | no | no | respiratory |
| ***B. lata*** **VC19230** | no | no | no | no | throat |
| ***B. metallica*** **VC8135** | no | no | no | yes | sputum |
| ***B. multivorans*** **VC12152** | no | no | yes | no | respiratory |
| ***B. multivorans*** **VC12539** | no | no | no | yes | respiratory |
| ***B. multivorans*** **VC12675** | no | no | no | no | sputum |
| ***B. multivorans*** **VC13125** | no | no | no | no | respiratory |
| ***B. multivorans*** **VC13145** | no | no | no | no | sputum |
| ***B. multivorans*** **VC14443** | no | no | no | no | sputum |
| ***B. multivorans*** **VC14749** | no | no | no | yes | sputum |
| ***B. multivorans*** **VC14757** | no | no | no | yes | respiratory |
| ***B. multivorans*** **VC15814** | no | no | no | no | respiratory |
| ***B. multivorans*** **VC16959** | no | no | no | no | sputum |
| ***B. multivorans*** **VC3419** | no | no | yes | yes | sputum |
| ***B. multivorans*** **VC4282** | no | no | no | yes | sputum |
| ***B. multivorans*** **VC6564** | no | no | yes | yes | sputum |
| ***B. multivorans*** **VC7102** | no | no | yes | yes | throat |
| ***B. multivorans*** **VC7704** | no | no | yes | yes | sputum |
| ***B. multivorans*** **VC7960** | no | no | no | no | respiratory |
| ***B. multivorans*** **VC9159** | no | no | no | no | respiratory |
| ***B. spp*** **VC14128** | no | no | no | no | respiratory |
| ***B. stabilis*** **VC6482** | no | no | yes | yes | sputum |
| ***B. stabilis*** **VC9042** | no | no | no | no | respiratory |
| ***B. vietnamiensis*** **VC10362** | no | no | no | no | respiratory |
| ***B. vietnamiensis*** **VC11253** | no | no | no | no | sputum |
| ***B. vietnamiensis*** **VC11275** | no | no | no | yes | respiratory |
| ***B. vietnamiensis*** **VC12002** | no | no | no | no | respiratory |
| ***B. vietnamiensis*** **VC15208** | no | no | no | no | sputum |
| ***B. vietnamiensis*** **VC15774** | no | no | no | yes | sputum |
| ***B. vietnamiensis*** **VC16431** | no | no | no | no | respiratory |
| ***B. vietnamiensis*** **VC18210** | no | no | no | no | respiratory |
| ***B. vietnamiensis*** **VC2824** | no | no | yes | yes | sputum |
| ***B. vietnamiensis*** **VC9237** | yes | no | yes | yes | respiratory |
| ***B. vietnamiensis*** **VC9752** | no | no | no | no | respiratory |

**Table S8. HMAQ production in different media for strains having the *hmqABCDEFG* operon in their genome.**

| Strains | Type | *hmq* operon | | HMAQs production | | |
|---|---|---|---|---|---|---|
| | | *hmqA* | *hmqG* | TSB | ASM | TSA |
| ***B. ambifaria*** **AMMD** | Environmental | + | + | - | + | + |
| ***B. ambifaria*** **AU0212** | Clinical | + | + | + | + | + |
| ***B. ambifaria*** **AU4157** | Clinical | + | + | + | + | - |
| ***B. ambifaria*** **AU7994** | Clinical | + | + | - | - | - |
| ***B. ambifaria*** **CEP0617** | Clinical | + | + | + | + | - |
| ***B. ambifaria*** **CEP0958** | Clinical | + | + | + | + | + |
| ***B. ambifaria*** **CEP0990** | Clinical | + | + | + | - | - |
| ***B. ambifaria*** **CEP0996** | Clinical | + | + | + | + | + |
| ***B. ambifaria*** **CEP1231** | Clinical | + | + | - | + | - |
| ***B. ambifaria*** **ES0020** | Environmental | + | + | + | + | + |
| ***B. ambifaria*** **HI2425** | Environmental | + | + | - | + | + |
| ***B. ambifaria*** **HI2468** | Environmental | + | + | + | + | + |
| ***B. ambifaria*** **HI2482** | Environmental | + | + | + | + | + |
| ***B. ambifaria*** **HI2626** | Environmental | + | + | + | + | + |
| ***B. ambifaria*** **HI2672** | Environmental | + | + | + | + | + |
| ***B. ambifaria*** **HI3709** | Environmental | + | + | + | + | + |
| ***B. ambifaria*** **HI3738** | Environmental | + | + | + | + | + |
| ***B. ambifaria*** **HSJ1** | Clinical | + | + | + | + | + |
| ***B. ambifaria*** **PC736** | Environmental | + | + | + | + | + |
| ***B. ambifaria*** **VC11631** | Clinical | + | + | - | - | - |
| ***B. ambifaria*** **VC15422** | Clinical | + | + | + | + | + |
| ***B. ambifaria*** **VC16196** | Clinical | + | + | + | + | + |
| ***B. anthina*** **VC15382** | Clinical | + | + | - | - | - |
| **B. cepacia BTS13** | Clinical | + | + | + | - | - |
| **B. cepacia HI2430** | Environmental | + | + | + | + | + |
| **B. cepacia HI2563** | Environmental | + | + | + | + | + |
| **B. cepacia HI2578** | Environmental | + | + | + | + | + |
| **B. cepacia HI2615** | Environmental | + | + | + | + | + |
| **B. cepacia HI2671** | Environmental | + | + | + | + | + |
| **B. cepacia HI2741** | Environmental | + | + | + | + | + |
| **B. cepacia HI3312** | Environmental | + | + | + | + | + |
| **B. cepacia HI3551** | Environmental | + | + | + | + | + |
| **B. cepacia HI3708** | Environmental | + | + | + | + | + |
| **B. cepacia HI3895** | Environmental | + | + | + | + | + |
| **B. cepacia HI4577** | Environmental | + | + | + | - | + |
| **B. cepacia ATCC25416** | Environmental | + | + | - | - | - |
| **B. cepacia VC13132** | Clinical | + | + | + | + | + |
| **B. cepacia VC13196** | Clinical | + | + | - | - | - |
| **B. cepacia VC13394** | Clinical | + | + | + | + | + |
| **B. cepacia VC13575** | Clinical | + | + | - | - | - |
| **B. cepacia VC14106** | Clinical | + | + | + | + | + |
| **B. cepacia VC14457** | Clinical | + | + | + | + | + |
| **B. cepacia VC17333** | Clinical | + | + | + | + | + |
| **B. cepacia VC17746** | Clinical | + | + | + | + | + |
| **B. cepacia VC17928** | Clinical | + | + | + | + | + |
| **B. cepacia VC18315** | Clinical | + | + | + | + | + |
| **B. cepacia VC18839** | Clinical | + | + | + | + | + |
| **B. cepacia VC18842** | Clinical | + | + | + | + | + |
| **B. cepacia VC19225** | Clinical | + | + | - | + | + |
| **B. cepacia VC19276** | Clinical | + | + | + | + | + |
| **B. cepacia VC9490** | Clinical | + | + | + | + | + |
| **B. contaminans FFH2055** | Clinical | + | + | + | - | - |
| **B. contaminans HI3570** | Environmental | + | + | + | + | + |
| **B. contaminans HI3852** | Environmental | + | + | + | + | + |
| **B. contaminans HI4067** | Environmental | + | + | + | + | + |

| | | | | | | |
|---|---|---|---|---|---|---|
| **B. contaminans HI4232** | Environmental | + | + | + | + | + |
| **B. contaminans VC14347** | Clinical | + | + | + | + | + |
| **B. contaminans VC15406** | Clinical | + | + | - | - | + |
| **B. contaminans VC16087** | Clinical | + | + | + | + | + |
| **B. contaminans VC16848-b** | Clinical | + | + | + | + | + |
| **B. contaminans VC16897** | Clinical | + | + | - | - | - |
| **B. contaminans VC16948** | Clinical | + | + | - | - | + |
| **B. contaminans VC19056** | Clinical | + | + | + | + | + |
| **B. contaminans VC19124** | Clinical | + | + | + | + | + |
| **B. contaminans VC9624** | Clinical | + | + | + | + | + |
| **B. diffusa VC14008** | Clinical | + | + | - | - | - |
| **B. dolosa LMG21443** | Environmental | + | + | - | - | - |
| **B. lata BC01** | Environmental | + | + | + | + | + |
| **B. lata VC6377** | Clinical | + | + | - | + | + |
| **B. metallica ES0559** | Environmental | + | + | - | - | - |
| **B. pyrrocinia Bcc indeterminate 2 BC02** | Environmental | + | + | + | + | - |
| **B. pyrrocinia Bcc indeterminate 5 HI2575** | Environmental | + | + | + | + | + |
| **B. pyrrocinia Bcc indeterminate 5 HI2690** | Environmental | + | + | + | + | + |
| **B. pyrrocinia Bcc indeterminate 5 HI2701** | Environmental | + | + | + | + | + |
| **B. pyrrocinia Bcc indeterminate 5 HI3892** | Environmental | + | + | + | + | + |
| **B. pyrrocinia Bcc indeterminate 9 ES0209** | Environmental | + | + | - | + | + |
| **B. pyrrocinia CH-67 (LMG14191)** | Environmental | + | + | + | + | + |
| **B. seminalis HI2490** | Environmental | + | + | - | + | - |
| **B. stagnalis Bcc indeterminate 6 HI3537** | Environmental | + | + | - | + | + |
| **B. stagnalis HI2720** | Environmental | + | + | - | + | - |
| **B. stagnalisMSMB1956WGS** | Environmental | + | + | + | + | - |
| **B. territorii MSMB1301WGS** | Environmental | + | + | + | + | + |
| **B. territorii MSMB1502WGS** | Environmental | + | + | + | + | + |
| **B. ubonensis LMG24263** | Clinical | + | + | - | - | - |
| **B. vietnamiensis CEP0040** | Clinical | + | + | - | - | - |
| **B. vietnamiensis HI3392** | Environmental | + | + | - | + | + |
| **B. vietnamiensis VC17180** | Clinical | + | + | - | - | - |
| **B. vietnamiensis VC8245** | Clinical | + | + | + | + | + |
| **B. vietnamiensis VC9237** | Clinical | + | + | - | - | - |
| **other Bcc Bcc indeterminate 1 BC06** | Environmental | + | + | + | + | + |
| **other Bcc Bcc indeterminate 3 BC13** | Environmental | + | + | + | + | + |
| **other Bcc Bcc indeterminate 3 ES0139** | Environmental | + | + | + | + | + |
| **other Bcc Bcc indeterminate 4 BC04** | Environmental | + | + | + | + | + |
| **other Bcc Bcc indeterminate 5 BC03** | Environmental | + | + | + | + | + |

**Table S9. Primers used in this study.**

| Name | Sequence | Function | Reference |
|---|---|---|---|
| hisA_Bcc_F | GGTCGACCTGAACGGCGC | Reference gene for PCR screening | Papaleo et al. 2010 |
| hisA_Bcc_R | CGTCGGTCGCGACCTTGCC | | Papaleo et al. 2010 |
| hmqA_Bcc_F | CCGCTCGCGTTYACGTTYGG | Amplification of *hmqA* gene in Bcc - degenerated | This study |
| hmqA_Bcc_R | CCCGTCAGGTTCCAGCCG | | This study |
| hmqG_Bcc_F | GGCGTCGCAGGAAATCACG | Amplification of *hmqG* gene in Bcc - degenerated | This study |
| hmqG_Bcc_R | CGCGACACGAARTGCATGCC | | This study |
| hmqA_semirandom_3F | CCGTAAACAGGATGCTGACC | Semi random PCR round 1 | This study |
| CERK2A | GGCCACGCGTCGACTAGTACNNNNNNNNNNAGAG | | Jacobs et al. 2008 |
| CERK2B | GGCCACGCGTCGACTAGTACNNNNNNNNNNACGCC | | Jacobs et al. 2008 |
| CERK2C | GGCCACGCGTCGACTAGTACNNNNNNNNNNGATAT | | Jacobs et al. 2008 |
| hmqA_semirandom_2F | CTTTCGGCAACAGCACGAC | Semi random PCR round 2 | This study |
| CERK4 | GGCCACGCGTCGACTAGTAC | Semi random PCR round 2 | Jacobs et al. 2008 |
| hmqA_semirandom_F | TGATCGGAAACAGCAGCAGC | Sanger sequencing 5' hmqA gene | This study |
| recA_Bcc_F | GATAGCAAGAAGGGCTCC | Identification of Bcc | Pubmlst |
| recA_Bcc_R | CTCTTCTTCGTCCATCGCCTC | | Pubmlst |
| gyrB_Bcc_F | CGACAACTCGATCGACGA | | Pubmlst |
| gyrB_Bcc_R | GACAGCAGCTTGTCGTAG | | Pubmlst |
| recA_Bcc_seq_F | TGACCGCCGAGAAGAGCAA | Sanger sequencing | Pubmlst |
| recA_Bcc__seqR | GACCGAGTCGATGACGAT | | Pubmlst |
| gyrB_Bcc_seq_F | ATCGTGATGACCGAGCTG | | Pubmlst |
| gyrB_Bcc__seqR | CGTTGTAGCTGTCGTTCC | | Pubmlst |
| ndh_RTPCR_F | GCTCGGCTACGACGATCT | Reference gene for RT-PCR | This study |
| ndh_RTPCR_R | GGCCTGGTCGAGGTTTTC | | This study |
| hmqA_Bambi_RTPCR_F | CTTGCCCCCTGCCGAAGATT | Expression of *hmqA* by RT-PCR | This study |
| hmqA_Bambi_RTPCR_R | CGGCGCAATTGAGAAACGG | | This study |
| hmqA_Bcep_RTPCR_F | CGACTTTTGCCGCGCGAA | | This study |
| hmqA_Bcep_RTPCR_R | GCAATTGAGAAACGGCGG | | This study |
| hmqA_Bcont_RTPCR_F | TTTTGCCGCGCGAACCTG | | This study |
| hmqA_Bcont_RTPCR_R | TTGAGAAACGGCGGGAACTG | | This study |
| hmqA_Bviet_RTPCR_F | TTTGCCCCCTGCCGAGGATT | | This study |
| hmqA_Bviet_RTPCR_R | CGGCGCAATTGAGAAACGG | | This study |

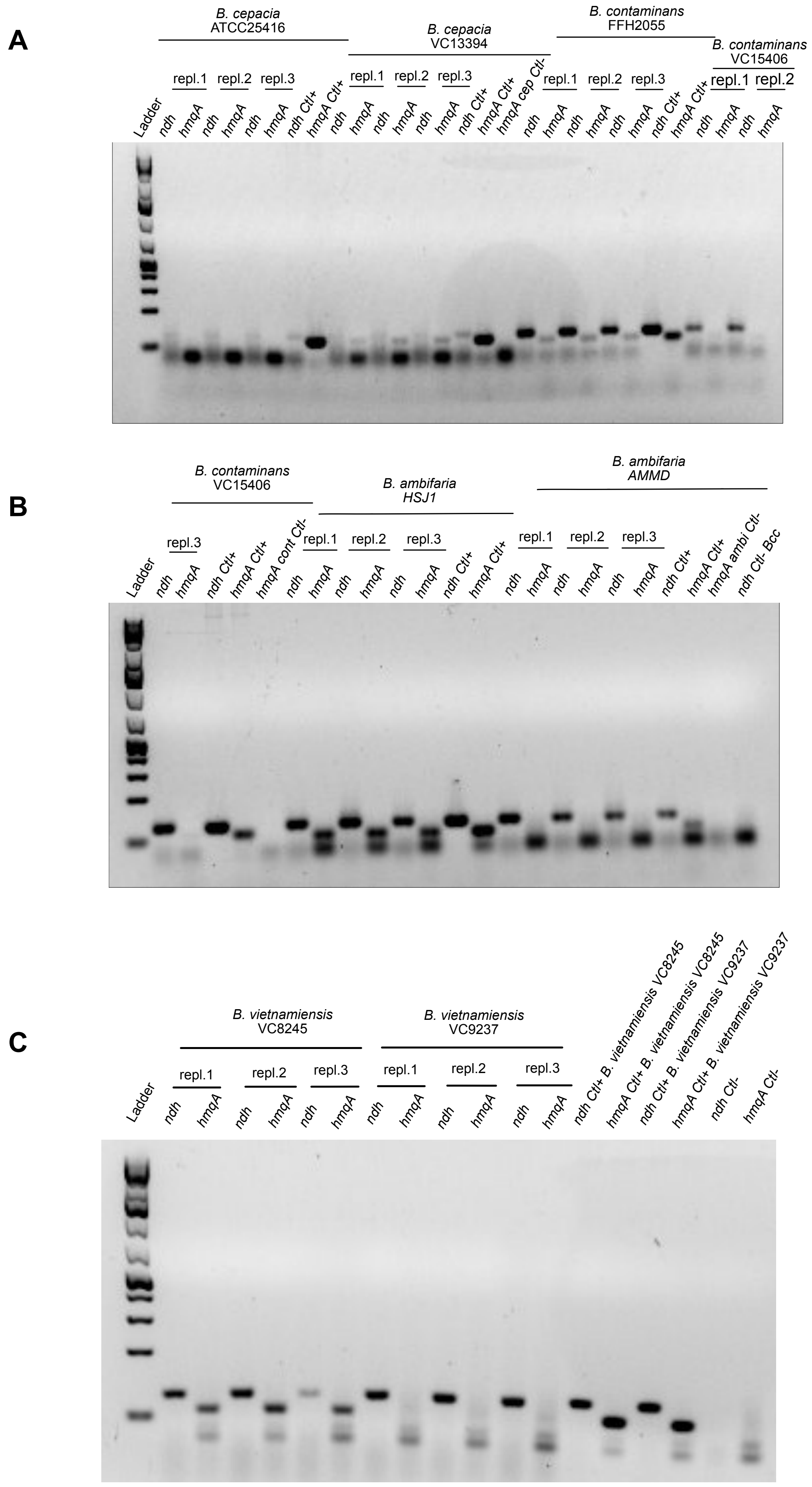
A
B. cepacia
ATCC25416
B. cepacia
VC13394
B. contaminans
FFH2055
B. contaminans
VC15406
repl.1
repl.2
repl.3
Ladder
ndh
hmqA
ndh Ctl+
hmqA Ctl+
hmqA cep Ctl-
B
B. contaminans
VC15406
B. ambifaria
HSJ1
B. ambifaria
AMMD
hmqA cont Ctl-
hmqA ambi Ctl-
ndh Ctl- Bcc
C
B. vietnamiensis
VC8245
B. vietnamiensis
VC9237
ndh Ctl+ B. vietnamiensis VC8245
hmqA Ctl+ B. vietnamiensis VC8245
ndh Ctl+ B. vietnamiensis VC9237
hmqA Ctl+ B. vietnamiensis VC9237
ndh Ctl-
hmqA Ctl-

**Figure S1. The expression of the *hmqA* gene in the main species of Bcc strains having the *hmqABCDEFG* operon but which do not produce HMAQs when cultured in TSB.** A) RT-PCR on B. *cepacia* and *B. contaminans* strains. B) RT-PCR on *B. contaminans* and *B. ambifaria* strains. C) RT-PCR on *B. vietnamiensis* strains.
